# Light-responsive monobodies for dynamic control of customizable protein binding

**DOI:** 10.1101/2020.03.08.831909

**Authors:** César Carrasco-López, Evan M. Zhao, Agnieszka A. Gil, Nathan Alam, Jared E. Toettcher, José L. Avalos

## Abstract

Customizable, high affinity protein-protein interactions, such as those mediated by antibodies and antibody-like molecules, are invaluable to basic and applied research and have become pillars for modern therapeutics. The ability to reversibly control the binding activity of these proteins to their targets on demand would significantly expand their applications in biotechnology, medicine, and research. Here we present, as proof-of-principle, a light-controlled monobody (OptoMB) that works *in vitro* and *in vivo*, whose affinity for its SH2-domain target exhibits a 300-fold shift in binding affinity upon illumination. We demonstrate that our αSH2-OptoMB can be used to purify SH2-tagged proteins directly from crude *E. coli* extract, achieving 99.8% purity and over 40% yield in a single purification step. This OptoMB belongs to a new class of light-sensitive protein binders we call OptoBinders (OptoBNDRs) which, by virtue of their ability to be designed to bind any protein of interest, have the potential to find new powerful applications as light-switchable binders of untagged proteins with high affinity and selectivity, and with the temporal and spatial precision afforded by light.

## Introduction

The high binding affinity and selectivity of antibodies and antibody-like proteins, such as monobodies, nanobodies, affibodies, anticalins and DARPins, have made them central tools of modern medicine, biotechnology, and basic and applied research^1–7^. Because of their ability to be raised, selected, or designed to bind practically any protein of interest^6,8,9^, these protein binders have found multiple applications in diagnostics^3,10,11^, therapeutics^4,12,13^ and biologics manufacturing^14–16^. As such, they have revolutionized the way we treat disease^13,17–19^, purify proteins^20–23^, and study biological phenomena^9,24–27^. The enormous impact that high affinity protein binders have had in medicine, biotechnology, and research stems from their usually irreversible binding to very specific targets. However, their repertoire of applications could still be greatly expanded if they were engineered with optogenetic control over their binding affinities, such that they could bind their targets instantly and reversibly depending on light conditions, while maintaining their characteristically high affinity and selectivity.

Protein binders can be developed from a variety of scaffolds including based on immunoglobulin domains^28,29^ or other small single-domain proteins^2,17^. Monobodies belong to a class of high affinity protein binders that have non-immunoglobulin scaffolds^30,31^. They are derived from the 10^th^ domain of human fibronectin, engineered to structurally and functionally resemble nanobodies^28,30,31^. As antibody mimetics with human origins, their use as biologic drugs is expected to substantially reduce unwanted immune responses^17,32^. Their high affinity, specificity and straightforward expression in multiple cell types also make them a versatile tool for research^6,31^. The small size and relative stability of monobodies (less than 100 amino acids), as well as their lack of disulfide bridges, robust activity inside^33^ and outside of cells^34^, and their ability to be selected to bind countless different proteins with high affinity and selectivity, make them ideal candidates to develop light-switchable protein binders by fusing them to light-responsive proteins.

A protein domain widely used for optogenetic tool development is the second light, oxygen, and voltage (LOV) domain from the oat, *Avena sativa*, photosensor Phototrophin 1, called the AsLOV2 domain^35^. This domain elicits its light response through a large conformational change (Fig. 1a) triggered by the formation of a covalent bond between a photoexcited flavin chromophore, FMN, and a conserved cysteine^36,37^. The C-terminal helix, Jα, of AsLOV2 is packed against its core domain in the dark. Upon blue light stimulation (optimally 447nm) the covalent bond between FMN and the conserved cysteine causes the Jαhelix to undock, become disordered and move away from the core domain ^38 39,40^. Back in the dark, the covalent bond with FMN decays, allowing the Jαhelix to fold back into its tightly packed dark state conformation^39,40^.

**Fig. 1:**
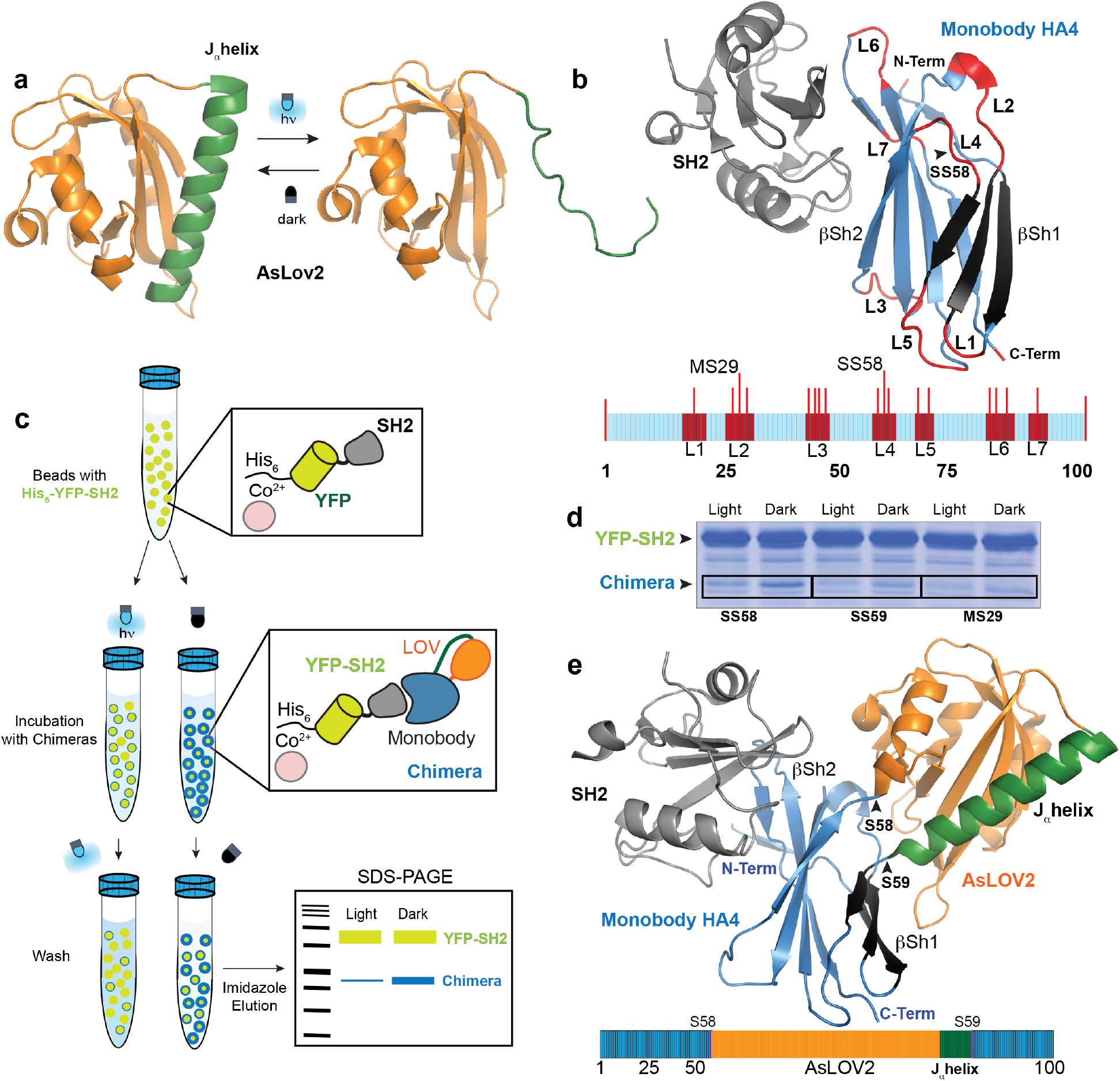
Design and screens of AsLOV2-Monobody chimeras and final OptoMB. **a**, Light-triggered conformational change of the Jαhelix (green) of AsLOV2. **b**, Crystal structure of Monobody HA4 (blue) bound to SH2 domain (gray). The monobody fold consists of two β-sheets, βSh1(black) and βSh2 (light blue), and seven structurally conserved loops (red), where AsLOV2 was inserted in our chimeras. Loop L4, where AsLOV2 is inserted in OptoMB, is shown with an arrow. The cartoon below shows the relative size and location of loops (red), including positions of AsLOV2 insertion (red lines) and insertion sites of identified light-responsive chimeras: MS29 and SS58 (in OptoMB). **c**, Schematic diagram of pull-down screens used to identify light-responsive chimeras. Co^2+^ agarose beads (pink) were used to immobilize His_6_-YFP-SH2 (yellow-gray), which were then incubated with HA4-AsLOV2 chimeras (blue-orange) in either dark or light. After washing and eluting with imidazole, the eluents were resolved on SDS-PAGE, were differences in protein band intensity between samples exposed to different light conditions reflect differences in chimera SH2 binding. **d**, Chimera pull-down screen results showing SDS-PAGE protein bands of two chimeras that bind better in the dark than in the light. These light-responsive chimeras have AsLOV2 inserted in either Loop L2 (MS29) or Loop L4 (SS58, SS59 and MS29). **e**, Energy-minimized structural model of the dark conformation of OptoMB with AsLOV2 (orange with the Jαhelix in green) inserted in position SS58 of HA4 (blue and black), interacting with the SH2 domain (gray).

Several systems have exploited this large, light-triggered, conformational change of AsLOV2 to confer light-dependence upon natural functions of different proteins. Insertion of AsLOV2 into solvent-exposed loops of kinases, phosphatases and guanine exchange factors make their enzymatic activities and downstream signaling events light-dependent^41^. A light-switchable Cas9 nuclease has also been developed using the AsLOV2 homolog from *Rhodobacter sphaeroides*, RsLOV, in which the light-triggered conformational change of RsLOV dissociates a sterically occluded RsLOV-Cas9 chimera homodimer to release Cas9 nuclease activity^42^. This approach was also demonstrated in smaller proteins by fusing AsLOV2 to the small natural CRISPR inhibitor AcrIIA4, resulting in a light-responsive anti-CRISPR system to control genome editing with light^43^. The versatility of AsLOV2 to bestow light-dependent functionalities to chimeric proteins in a variety of contexts, shown in these studies, encouraged us to use this light-responsive protein to develop light controls in a protein binder.

Here we show that by fusing a monobody to the AsLOV2 domain we obtain a light-dependent monobody, or OptoMonobody (OptoMB), whose binding affinity to the monobody’s cognate protein target is controllable with light. Taking a structure-based protein engineering approach, we inserted the AsLOV2 domain into structurally conserved, solvent exposed, loops of a monobody that binds the SH2 domain of Abl kinase. We show that one of these chimeras preferentially binds to the SH2 domain in the dark, both *in vitro* and in mammalian cells. We harness the ~300-fold change in binding affinity between lit and dark states to implement light-based protein purification^44^ using an OptoMB resin in what we call “light-controlled affinity chromatography” (LCAC). This work, and an accompanying study reporting light-dependent nanobodies or OptoNanobodies (OptoNBs)^45^, represent the first demonstrations of protein binders with high affinity and selectivity, engineered with light-control of their binding activity.

The new class of light-responsive protein binders, or OptoBinders (OptoBNDRs), which in principle can be engineered, screened, or selected to bind any protein of interest, have great potential to be used in numerous new applications in biotechnology, synthetic biology, and basic research.

## Results

### Design and selection of OptoMonobodies

To demonstrate the feasibility of developing a light-sensitive monobody, we chose the HA4 monobody, which binds with high affinity (*K_d_* ~7nM) to the SH2 domain of the human Abl kinase, *in vitro* and *in vivo*^46^. This is an interesting and valuable target, as many proteins containing SH2 domains in general, and Abl kinase in particular, are involved in human health and disease^47–49^. In addition, the availability of the crystal structure of HA4 bound to SH2 domain^46^ is a valuable resource for our rational protein engineering approach. Our strategy to develop a light-sensitive HA4 was to design various chimeras of this monobody with the light-oxygen-voltage-(LOV) sensing domain of Phototropin1 from *Avena sativa*, AsLOV2, and test their ability to bind and release the SH2 domain depending on light conditions.

To build our chimeras, we used a truncated version of AsLOV2, which induces light-dependent conformational changes in engineered nanobody chimeras more efficiently than its full-length counterpart^45^. Our strategy was to insert the shortened AsLOV2 domain in all seven structurally-conserved, solvent-exposed loops of HA4 (Fig. 1b). Given the large conformational change of AsLOV2 triggered by light (Fig. 1a), our hypothesis was that the native conformation of the monobody domain in some chimeras would be preserved in the dark, allowing it to bind to SH2, but disrupted in the light, causing it to release its target. Guided by the crystal structure of HA4 bound to SH2 (PDB ID: 3k2m)^46^, we explored potential sites within the seven solvent-exposed loops in HA4 where we could insert AsLOV2. We selected as many positions as possible in each loop, avoiding those were we have reasons to believe the dark state conformation of AsLOV2 would disrupt the core β-sheets of the monobody or interfere with the monobody-SH2 binding interaction. We also excluded positions where the light-triggered conformational change of the AsLOV2 Jαhelix might be impeded by clashes with the monobody core. After this structural analysis, we selected 17 AsLOV2 insertion sites across all solvent exposed loops of HA4, as well as N- or C-terminal fusions (Fig. 1b and Supplementary Table 1).

To find chimeras that can bind the SH2 domain in the dark but not in the light, we screened our constructs using an *in vitro* pull-down assay (Fig. 1c). First, we produced an N-terminally His-tagged fusion of yellow fluorescent protein (YFP) and SH2 domain (His_6_-YFP-SH2) in *E. coli*, and immobilized it onto cobalt-charged agarose beads. We then incubated the beads with crude extracts of *E. coli* expressing each of the different AsLOV2-HA4 chimeras, in either blue light or darkness. After washing the beads under the same light conditions (see Methods), we eluted with imidazole and resolved the products with denaturing polyacrylamide gel electrophoresis (SDS-PAGE) to analyze the binding of each chimera in different light conditions (Fig. 1c,d). We anticipated that chimeras that bind to SH2 preferentially in the dark would show a more intense band on SDS-PAGE for samples that were incubated and washed in the dark, relative to the samples treated in the light (Fig. 1c,d and Supplementary Fig. 1).

We found that AsLOV2 insertions in two different HA4 loops produce chimeras with the expected behavior in our pull-down assays. One promising chimera has AsLOV2 inserted between residues Met29 and Ser30 (a site we call MS29, following a naming system for sites used in this study), located in loop L2 of HA4 (Fig. 1b,d and Supplementary Fig. 1). We only saw an effect involving this loop when AsLOV2 was inserted at position MS29 and loop L2 was shortened by removing the three surrounding amino acids (Ser30, Ser31 and Ser32, see Supplementary Sequences). Another chimera with positive results has AsLOV2 inserted between Ser58 and Ser59 (site SS58) located in loop L4 of HA4 (Fig. 1b,d). Insertions at other positions within loop L4 (^57^YSSS^60^) show smaller degrees of variation in band intensity between beads treated in the light versus in the dark (Fig. 1d and Supplementary Fig. 1). This suggests that loop L4 is a “hot spot” for favorable orientations between AsLOV2 and the monobody to produce light-responsive chimeras that switch between a conformation that allows target binding (in the dark) and one that promotes target dissociation (in the light). Compared to AsLOV2 insertions at other positions in loop L4, the insertion at SS58 shows the largest qualitative difference between the chimera bound to beads in different light conditions (Fig. 1d and Supplementary Fig. 1). Therefore, we chose this chimera to continue our study, naming it αSH2 OptoMonobody (henceforth called OptoMB in this study). A structural model of this OptoMB (Fig. 1e and Supplementary Movie 1) reveals that the orientation of AsLOV2 in this chimera (modeled in the dark state) is compatible with SH2 binding and suggests a possible mechanism by which a light-triggered conformational change of the Jαhelix may disrupt the HA4 monobody to disfavor SH2 binding (see Discussion).

### *In vitro* characterization of OptoMB

The *in vitro* light-dependent interaction of our OptoMB and SH2 domain (Fig. 2a) can be visualized by fluorescence microscopy using immobilized OptoMB. We immobilized His-tagged OptoMBs harboring a mutation in AsLOV2 (V416L) that extends the lifetime of the photoactivated state^50^ (OptoMB_V416L_) onto Ni-charged agarose beads. Immobilized parental HA4 monobodies were used in parallel as a control. The monobody-coated beads were then incubated with a fusion of yellow fluorescent protein and SH2 (YFP-SH2) in the dark until they reached equilibrium and imaged over time using confocal microscopy in the presence or absence of blue light (450 nm). For OptoMB-coated beads, light exposure induced a rapid decrease in YFP signal on the surface of the bead within one imaging frame (2 min), consistent with light-triggered dissociation of the SH2 domain (Fig. 2b upper panel and Supplementary Movie 2). In contrast, no fluorescence change was observed under identical conditions for the control HA4-coated beads (Fig. 2b lower panel and Supplementary Movie 2). Localized illumination could also be used to restrict SH2 unbinding to a single bead in a crowded field (Supplementary Fig. 2a). In this case, YFP fluorescence was rapidly and reversibly controlled for the illuminated bead but not a nearby un-illuminated bead (Supplementary Fig. 2b and Supplementary Movie 3). These results demonstrate that OptoMBs provide temporal, spatial, and reversible control over protein binding.

**Fig. 2:**
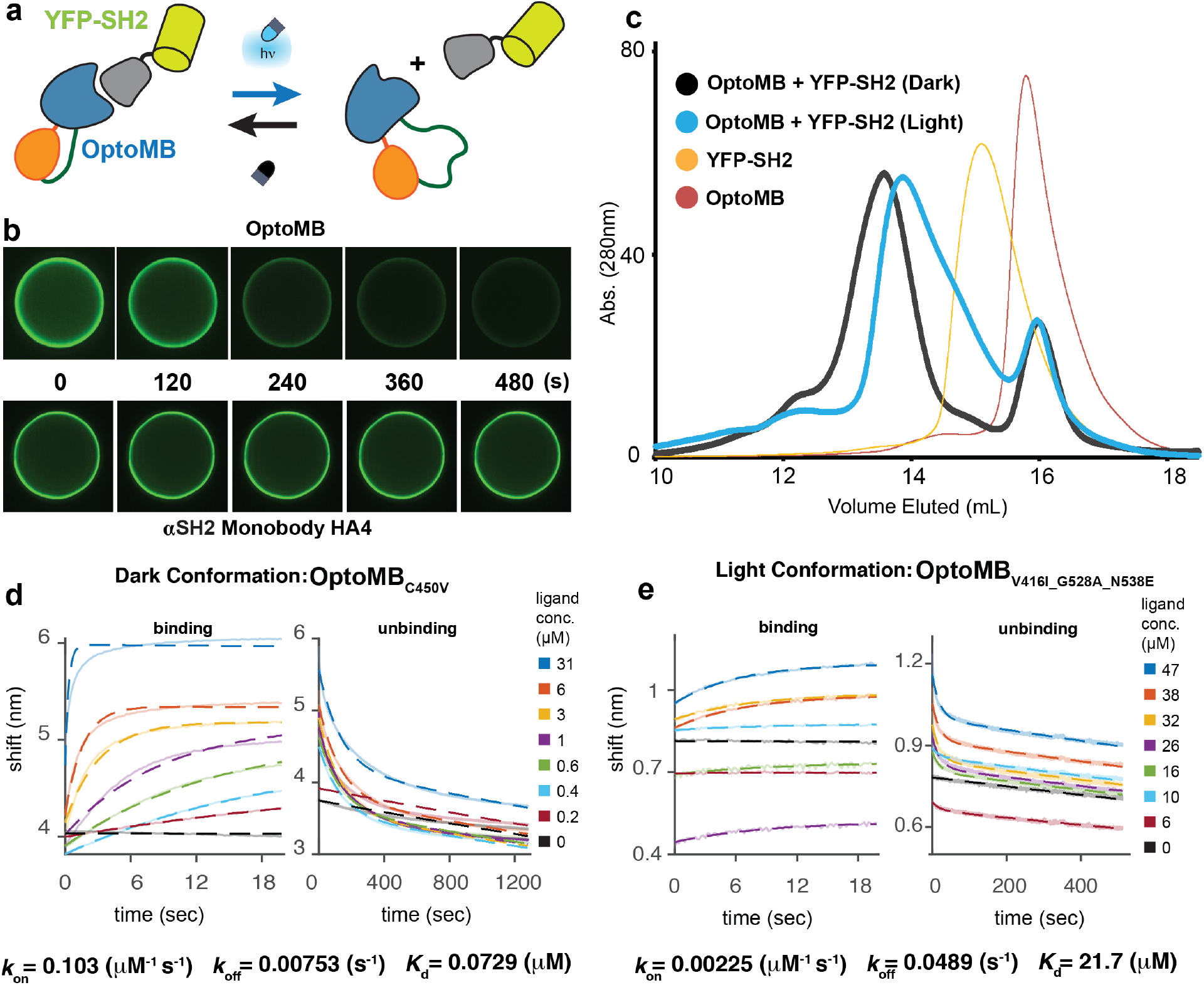
*In vitro* characterization of OptoMB. **a**, Schematic diagram of OptoMB and YFP-SH2 darkness-dependent interaction. In the dark OptoMB binds to SH2 domain of YFP-SH2 fusion; upon blue light stimulation, the AsLOV2 domain disrupts the HA4 domain of OptoMB, reducing its affinity to SH2. **b**, Time course of fluorescence microscopy images of YFP-SH2 binding to agarose beads conjugated with OptoMB_V416L_ (top panel) or HA4 as control (lower panel). Starting with beads incubated in the dark, time course begins upon blue light stimulation (t = 0), followed by a sequence of images taken every 120 s for a total of 480 seconds (see also Supplementary Movie X) **c**, Size exclusion chromatography profiles of OptoMB and YFP-SH2 interactions in light (blue line) and dark (black line). **d**, **e**, BLI measurements of binding (left) and unbinding (right) activities of YFP-SH2 to immobilized OptoMB in the dark, **d**, (using OptoMB_C450V_) or in the light, **e**, (using OptoMB_V416I_G528A_N538E_ fused to SUMO tag). The calculated rate and dissociation constants are shown below the BLI data.

The light dependency of OptoMB interactions with YFP-SH2 can also be analyzed in solution by size exclusion chromatography (SEC). We prepared mixtures of purified YFP-SH2 and OptoMB (or HA4 monobody, as control), in which OptoMB was added in excess (see Methods). We then loaded each sample to a gel filtration column under continuous darkness or blue light (Supplementary Fig. 2c). We found that for both the dark-incubated OptoMB sample (Fig. 2c) and the HA4 monobody in either light condition (Supplementary Fig. 2d), the YFP-SH2 elutes primarily as a monobody-bound complex. In contrast, the illuminated OptoMB sample shows a higher retention time for the YFP-SH2-OptoMB complex and a larger proportion of the YFP-SH2 eluting as a monomer on its own (Fig. 2c). Taken together, our bead-imaging and SEC data are both consistent with a blue light-triggered reduction in the OptoMB-SH2 binding affinity, both in solution and on protein-coated surfaces.

To quantify the changes in OptoMB-SH2 binding, we determined the kinetic rate constants and binding affinity in different light conditions. In these assays we took advantage of three classes of mutations in AsLOV2 to vary properties of the OptoMB binding switch. Bio-layer interferometry uses visible light for measuring changes in binding, so we first prepared a light-insensitive OptoMB by introducing the well-characterized C450V mutation in AsLOV2 that renders it light-insensitive^36,51^ (OptoMB_C450V_). Conversely, we ensured that illumination could drive efficient conversion to the lit state by generating additional OptoMB variants with mutations that extend the lifetime of the lit state after illumination from ~80 to 821 or up to 4300 sec *in vitro* (AsLOV2 V416I^49^ or V416L^52^ respectively). Finally, we added the AsLOV2 G528A and N538E mutations (to make the triple mutant OptoMB_V416I_G528A_N538E_), which have been reported to stabilize the dark state conformation and potentially decrease leakiness in the absence of illumination^53,54^. We fit the resulting binding and dissociation data to a mass-action kinetic binding model (Fig. 2d,e and Supplementary Fig. 2e,f), allowing us to obtain estimates of the rate constants of binding (*k*_on_) and unbinding (*k_off_*) as well as the overall dissociation constant (*K_d_*) of the OptoMB-SH2 interaction in different light conditions (Supplementary Table 2).

We found that the binding affinity of OptoMB to SH2 changes dramatically when switching from dark to light conditions. The dissociation constant of the OptoMB-SH2 interaction in the dark is *K_d_* = 0.07 μM, comparable to that of the HA4 monobody. However, in the light it increases to *K_d_* = 11.2 μM or *K_d_* = 21.7 μM, depending on which lit-stabilized OptoMB variant we use. This amounts to a 160-310 fold-change in binding affinity upon switching light conditions, which explains the light-dependent behaviors observed in the bead-imaging and SEC experiments above. These changes in *K_d_* arise primarily from the decrease in the OptoMB-SH2 binding rate constant (*k_on_*) in the light, although we also observe an increase in the unbinding rate constant (*k_off_*) (Supplementary Table 2). These data are consistent with a light-induced change in the conformation of the OptoMB that disrupts the binding interface, substantially decreasing the likelihood of a productive association between the lit-state OptoMB and its target SH2. Moreover, the dominant role of *k_on_* variation in light-induced changes in binding affinity is consistent with our proposed OptoMB functional model, which opens the door to further engineering and optimization (see Discussion).

### Light-controlled affinity chromatography (LCAC) with immobilized OptoMB

We reasoned that the substantial change in OptoMB binding affinity could open the door to purifying a protein of interest simply by shifting illumination conditions (Fig. 3a), a procedure that we termed “light-controlled affinity chromatography” (LCAC). We immobilized two variants of His-tagged OptoMBs harboring either the wild-type AsLOV2 or the triple-mutant (OptoMB_V416I_G528A_N538E_) described above, onto Co-charged Agarose beads to make an αSH2-OptoMB affinity resin. We then incubated OptoMB-coated beads with crude lysate from *E. coli* overexpressing YFP-SH2. After washing thoroughly in the dark (see Methods), we eluted with blue light either in batch (Fig. 3b) or in a column (Supplementary Fig. 3). After elution, beads were washed with imidazole to recover any remaining protein bound to the beads in order to estimate the capacity and yields of the resin. With these initial LCAC purification trials, we achieved 95-98% purity in a single step, with yields ranging from 18-30% and binding capacities from 112-145 nmol (4.5-6 mg) of SH2-tagged YFP per mL of OptoMB resin, depending on the OptoMB variant used (Supplementary Table 3).

**Fig. 3:**
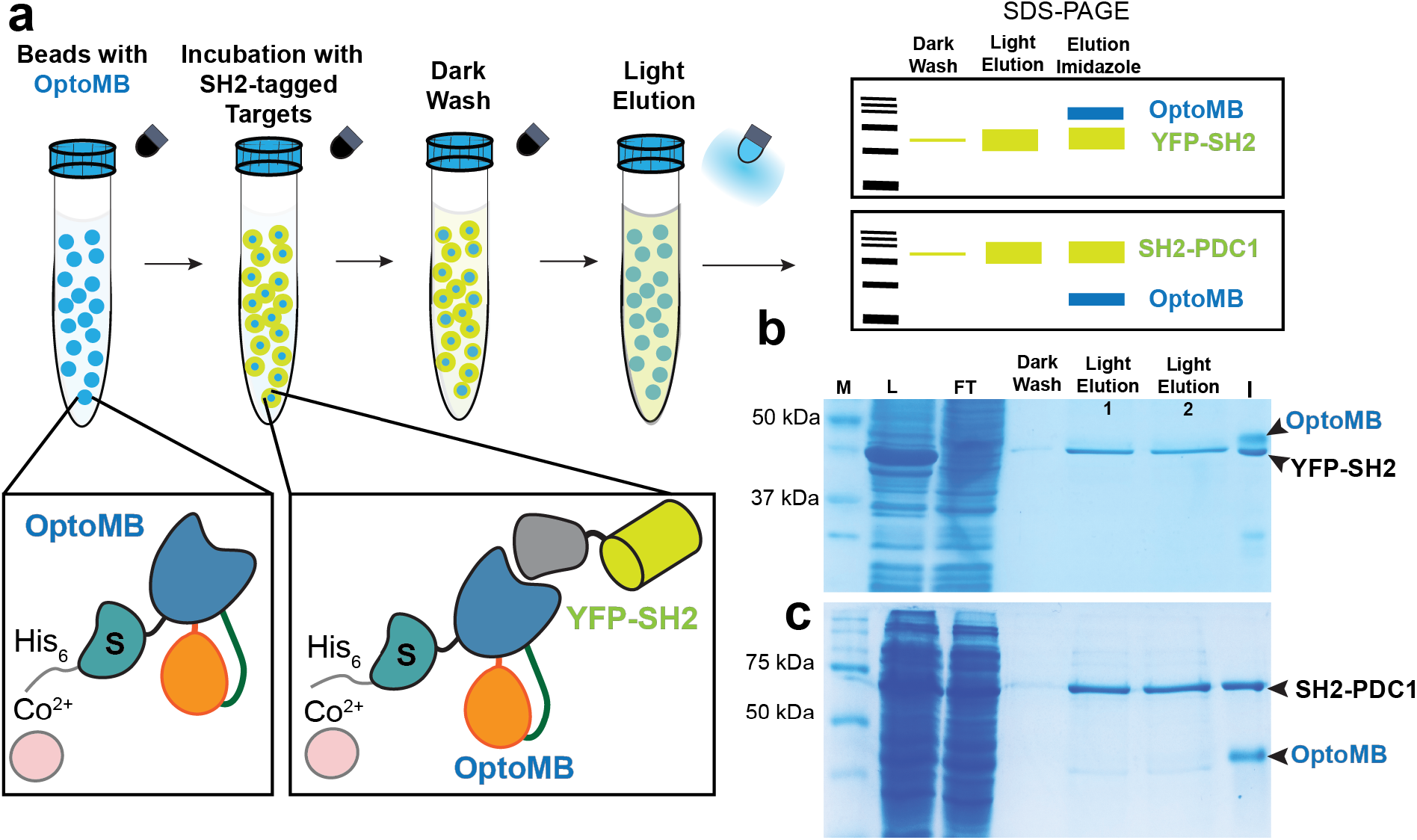
Light-Controlled Affinity Chromatography (LCAC) to purify SH2-tagged proteins using cobalt-immobilized OptoMB. **a**, Schematic diagram of functionalization of LCAC agarose beads using His_6_-OptoMB_V416I_G528A_N538E_ fused to an N-terminus SUMO tag (S) (bound to Co^2+^ beads) and LCAC procedure involving incubation of crude *E. coli* extract, and washing in the dark, followed by elution with blue light in batch or column (Supplementary Fig. 3). Finally, SDS-PAGE was used to resolve the fractions from each purification step. **b**, **c,** SDS PAGE gel of LCAC-purified YFP-SH2 (b), and SH2-PDC1 (c). Molecular weight markers (M), lysate (L), unbound flow through (FT), washing step in the dark (Dark Wash), two consecutives light elution aliquots (Light Elution 1 and 2), and the imidazole elution (I) were resolved in 12% SDS-PAGE.

To test whether LCAC could be applied to larger and more complex proteins, we also used it to purify the main pyruvate decarboxylase from *Saccharomyces cerevisiae*, Pdc1p. This enzyme catalyzes the decarboxylation of pyruvate to acetaldehyde for ethanol fermentation, and is composed of a homotetramer of 61 kDa monomers^55^, significantly larger than YFP. We fused the SH2 domain to the N-terminus of Pdc1p (SH2-PDC1) and performed LCAC to purify it from crude *E. coli* lysate, as described above, using a resin coated with OptoMB_V416I_G528A_N538E_. This procedure enabled purification of Pdc1p to 96% purity with a 39% yield (Fig. 3c and Supplementary Table 3). It is noteworthy that this purification works considerably well, despite the potential binding avidity of Pdc1p tetramers that would be predicted to increase the protein’s apparent affinity in both the light and dark. These results demonstrate that LCAC can be applied to purify relatively large proteins with quaternary structures of up to at least 300 kDa (including the fused SH2 domain), achieving a high degree of purity and an acceptable yield. Our results with YFP-SH2 and SH2-PDC1 further demonstrate that OptoMB-assisted purification is compatible with both N- and C-terminal SH2 tags.

While metal-affinity beads are effective at immobilizing OptoMB for LCAC (Fig. 3 and Supplementary Fig. 3), they may be incompatible with some protein purification methods. Thus, to determine whether an alternative resin could be used for LCAC, we immobilized our OptoMB_V416I_G528A_N538E_ onto cyanogen bromide-activated sepharose beads (CNBr-beads), which immobilizes proteins by making covalent bonds with its primary amines (see Methods). Following the same purification protocol as above, we found that CNBr-beads are also effective at purifying both YFP and Pdc1p (Supplementary Fig. 4). A single step of CNBr-based purification achieved yields above 40% and purity of 96.7–99.8%, surpassing any other LCAC method tested (Supplementary Table 3). These gains are likely related to the reduced non-specific binding of *E. coli* proteins to CNBr-sepharose and covalent OptoMB attachment which allowed for more extensive washing at higher salt concentrations. We observed that the total loading capacity is not as high as that of OptoMB-coated Co-agarose (Supplementary Table 3), probably because random crosslinking to the CNBr-beads inactivates a significant fraction of the OptoMB by occluding its binding surface to SH2. Although much could still be done to further improve LCAC by optimizing the resin, method, or amino acid sequence of the OptoMB, these experiments demonstrate the feasibility of a practical *in vitro* application of light-responsive monobodies for protein purification.

### Light-dependent OptoMB binding *in vivo*

We have shown that OptoMB-SH2 binding can be controlled with light *in vitro*; as a final test of this system, we assessed whether similar control can be achieved in live mammalian cells where monobodies have been reported to work^33^. We transduced HEK293T cells with lentiviral vectors encoding a membrane-localized, fluorescent SH2 target protein (SH2-mCherry-CAAX) and cytosolic fluorescent OptoMB (OptoMB-irFP), reasoning that a light-dependent change in SH2-OptoMB binding would cause the OptoMB to redistribute between the cytosol and plasma membrane (PM) (Fig. 4a), as has been observed for conventional optogenetic protein-protein interactions in previous studies^56–58^. As a control, we expressed irFP-labeled HA4 monobody instead of OptoMB, which would be expected to bind to the membrane-localized SH2-mCherry-CAAX regardless of illumination conditions.

**Fig. 4:**
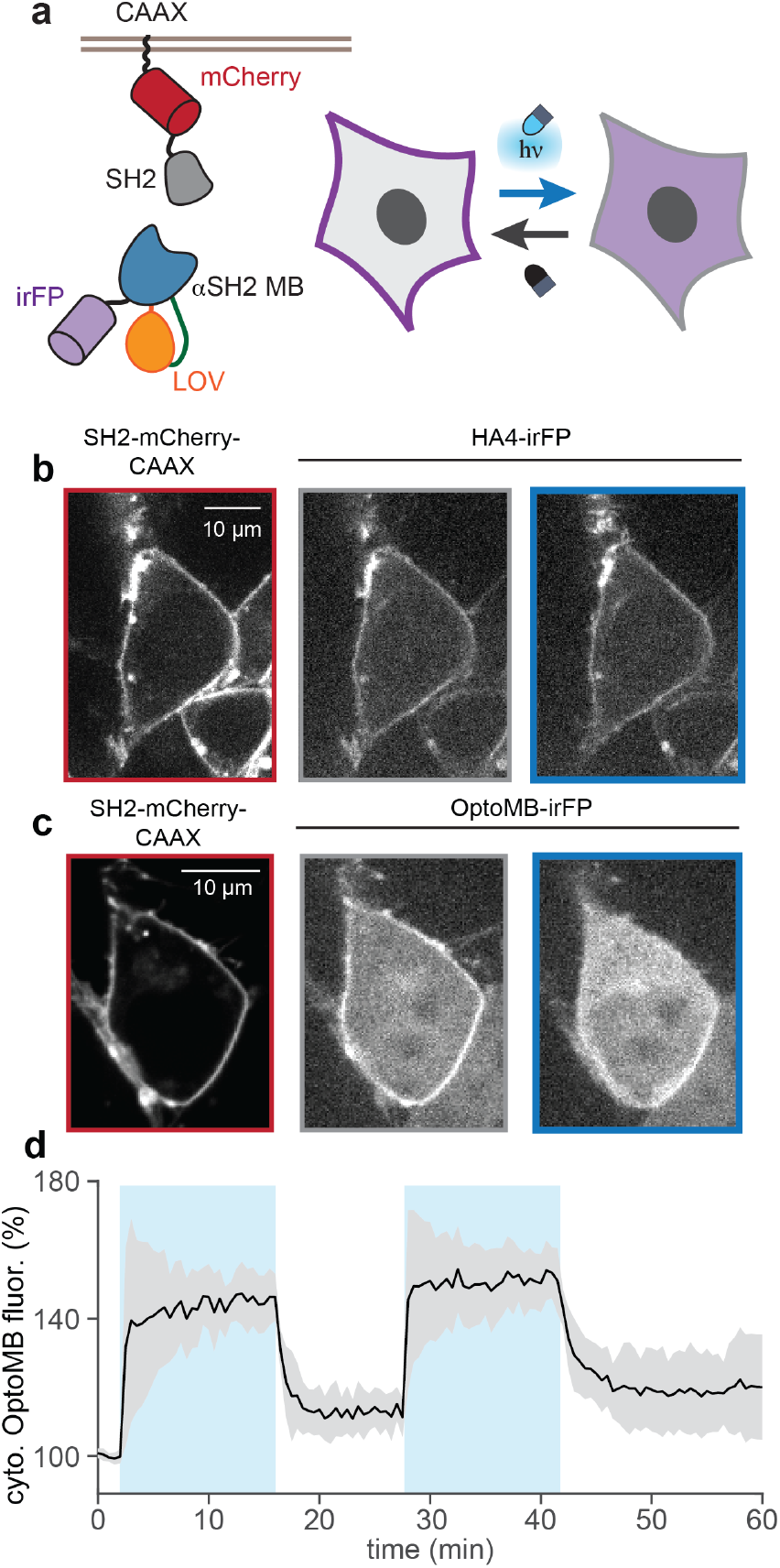
In vivo characterization of OptoMB using mammalian cells. **a**, Schematic diagram of the membrane-binding assay used, where irFP-tagged OptoMB (blue-orange-purple), binds to a fusion of SH2 (gray), mCherry (red), and CAAX (black linker), anchored to the plasma membrane (PM) of HEK 293T cells. In the dark, OptoMB-irFP binds to the membrane-bound SH2 enhancing irFP signal at the PM, reducing its cytosolic signal. In blue light, OptoMB releases SH2, causing a reduction in irFP signal at the membrane and an increase in the cytosol. **b**, HEK 293T cells expressing SH2-mCherry-CAAX and irFP-tagged HA4 monobody, as control, imaged in the dark or blue light. Left panel shows mCherry fluorescence of SH2 fusion anchored to the PM; central and right panels show HA4-irFP fluorescence in the dark and after 30 s of blue light stimulation, respectively, showing irFP-HA4 localized to the cytosol in either light condition. **c**, HEK 293T cells expressing SH2-mCherry-CAAX and OptoMB-iRFP imaged in the dark or blue light. Left panel, shows mCherry fluorescence of the SH2 fusion anchored to the PM. Central and right panels show HA4-irFP fluorescence in the dark and after 30 s of blue light stimulation, respectively, showing OptoMB enriched at the PM in the dark and in the cytosol in the light. **d**, Cytosolic irFP-HA4 fluorescence changing over time due to periodic pulses of blue light (blue sections). The fluorescence is expressed in percentage from the original value of cells exposed to the dark. Curve and shaded regions indicate mean ± SD for at least 10 cells respectively. White scale bar in the cell images is 10 μm.

Fluorescence imaging confirmed the PM localization of SH2-mCherry-CAAX (Fig. 4b,c left panels) as well as constitutively PM-bound HA4-irFP (Fig. 4b). In contrast, we observe a light-dependent shift in OptoMB localization (Fig. 4c), with PM enrichment in the dark and rapid redistribution to the cytosol upon light stimulation. Applying cycles of light and darkness further revealed that light-controlled binding is fully reversible intracellularly (Fig. 4d; Supplementary Movie 4). These experiments demonstrate functional photoswitching of OptoMB binding inside cells, opening the door to its *in vivo* application.

## Discussion

Here we show that by taking a rational protein engineering approach it is possible to develop a light-switchable monobody (OptoMB). Our OptoMB design strategy was based on the fusion of a light-switchable AsLOV2 domain to a structurally conserved loop of the H4A monobody which binds with high affinity and selectivity to the SH2 domain of the human Abl kinase. We find that this chimera binds to SH2 in the dark with similar affinity as its parental monobody H4A; but in blue light its binding affinity drops 150- to 300-fold relative to the dark state (Supplementary Table 2). Furthermore, the light responsiveness of OptoMB is effective to control binding to proteins fused to SH2 at either their N- or C-terminus. OptoMBs, along with the light-dependent nanobodies (OptoNBs) we describe in an accompanying study^45^, belong to a new class of light-dependent protein binders we call OptoBinders (OptoBNDRs), which offer promising new *in vivo* and *in vitro* applications.

A close inspection of the structural model of OptoMB and binding measurements suggest a possible mechanism for the light-dependent binding affinity of OptoMB. The monobody protein fold consists of two antiparallel β-sheets (βSh1 and βSh2) that interact with each other to form the protein core (Fig. 1b). In our original chimera screens, we inserted AsLOV2 in all intervening loops within (L1, L3, L6 and L7) and between (L2, L4 and L5) these β-sheets. The only chimeras that show a change in binding affinity in different light conditions are those with AsLOV2 inserted in loops L2 and L4, with L4 being the only loop with positive results at multiple insertion sites. Both of these loops connect βSh1 and βSh2 with each other, suggesting that chimeras involving either loop may act by a similar mechanism, where the light-triggered conformational change of the Jαhelix pulls the βSh1 and βSh2 apart from each other. Chimeras with AsLOV2 inserted in loop L5, located at the opposite side of the βSh1-βSh2 interaction relative to L2 and L4, show equally faint SDS-PAGE bands in either light condition, suggesting limited expression in *E. coli*, or weak binding to SH2 independently of light. The consequence of pulling on βSh1 and βSh2 from loops L2 or L4 is probably at least a partial disruption of the interactions and angle between them. This in turn would likely change the curvature of βSh2, which defines the shape of the paratope-like surface of H4A and its specific binding interactions with SH2^46^. Interestingly, because the conserved fold of monobodies always includes βSh1 and βSh2 interactions^31^, this light-triggered disruption of the binding surface may be transferable to other monobodies. It also suggests that mutating residues involved in βSh1-βSh2 interactions may provide some opportunities to tune the photoswitchable behavior of the OptoMB, by stabilizing either the dark- or lit-state conformations.

This model of light-induced disruption of the monobody’s target-binding site is also consistent with our measurements of binding kinetics. We originally reasoned that light stimulation might strain the OptoMB-SH2 interaction causing them to dissociate without necessarily inducing dramatic changes in binding site accessibility, thus predicting a light-induced change in the OptoMB-SH2 off-rate (*k_off_*). While we do observe as much as a 10-fold increase in *k_off_* in the light, consistent with this hypothesis, we found that most of the change in affinity comes from as much as a 46-fold change in the binding rate (*k_on_*) between dark and light conditions (Supplementary Table 2). This suggests that the binding site is disrupted enough to inhibit OptoMB-SH2 association in the first place. Yet despite what is likely to be a substantial conformational change in the OptoMB, we find that productive binding is fully reversible *in vitro* (Supplementary Fig. 2b and Supplementary Movie 3) and *in vivo* (Fig. 4d and Supplementary Movie 4). Overall, these data are consistent with a large change in the overall orientation or conformation of the βSh1-βSh2 sheets, disrupting binding but without driving irreversible protein misfolding. It is possible that the small size of the monobody domain, the short range of interactions between βSh1 and βSh2, and the lack of disulfide bonds in the monobody fold facilitate this high efficiency of interconversion between binding states.

Comparison of our binding data to a maximum theoretical value that assumes perfect transmission of energy between AsLOV2 and the monobody’s binding interface suggests that light-induced changes in OptoMB could be further improved. Previous studies have measured the free energy available from the dark-to-light conformational change of AsLOV2 using NMR spectroscopy^59^ and developed analytical models to study equilibrium constants of the dark- and lit-states of AsLOV2^53^. These studies predict that 3.8 kcal/mol of energy is transmitted from the absorption of one photon to structural rearrangements in AsLOV2. If all of this energy was transmitted to a change in OptoMB binding, it would result in a ~600-fold difference in binding affinity between the dark and lit states. Our BLI experiments measured a change in *K_d_* values of 150- to 300-fold, which is within the same order of magnitude of this maximum theoretical prediction, indicating that the light-induced conformational change of AsLOV2 is efficiently transmitted to disrupt the SH2 binding surface of the monobody domain. Nevertheless, it also suggests that further improvements might be achieved by optimizing the insertion site or linkers between the AsLOV2 and monobody, or by engineering the H4A domain to improve the light-induced allosteric coupling between the AsLOV2 domain and SH2-binding site. It is also possible that performance for particular applications might be improved by tuning the OptoMB-SH2 affinity to either weaker or tighter values. Finally, we note that the AsLOV2 domain might also be altered to tune the efficiency of the light-induced conformational change, for instance by further stabilizing the lit or dark state conformations or by altering its photoswitching kinetics (Fig. 3b, c and Supplementary Table 2). In principle, these changes could increase the overall energy beyond the 3.8 kcal/mol measured for the wild-type AsLOV2 domain. The structural targets set for optimization and the expected results will of course vary depending on the particular application objectives and the individual OptoMB/ligand pair in question.

The performance of our light-dependent OptoMB/SH2 binding pair is comparable to that of other optical dimerization systems previously used to control transcription, protein-protein interactions, or protein localization^60–62^. However, these previous systems have been developed by fusing proteins of interest to specific light-dependent interaction partners (such as PhyB/PIF3 or Cry2/CIB) evolved in photosensitive organisms. While the demonstrations we used in this study rely on fusing SH2 to different proteins, the true potential of this technology is not so much to replace the light-responsive tags used in previous optogenetic systems with SH2 and OptoMB, but in the possibility of engineering other monobodies against different targets of interest to make their specific interaction light-dependent. The relative simplicity of the monobody fold and the likely structural mechanism that confers light-dependent binding makes it reasonable to expect that other monobodies could be engineered to be light-dependent; starting from inserting AsLOV2 in loop L4 and then optimizing the precise position of the insertion site and the linkers. Monobodies could in principle be designed and selected to bind any protein of interest^6,31^ or to different epitopes on a single a target^31^. Our approach may thus be useful to design reversible interactions to a protein of interest without the need for a binding tag, which for some proteins may be impractical, not possible, or interfere with their natural activity.

## Methods

### Plasmid construction of chimeras for bacterial expression

One-step isothermal assembly reactions (Gibson assembly) were performed using previously described methods^61^. The monobody HA4 and the SH2 domain (codon optimized for *E. coli* expression) were ordered as gBlocks from Integrated DNA Technologies (IDT) containing homology arms. The following vectors from the pCri system^63^ were used: pCri-7b for constructs without a 6x-histidine tag; pCri-8b for constructs with a 6x-histidine tag (N-terminus), and pCri-11b for constructs with both SUMO and 6x-histidine tags (N-terminus). As the pCri vectors contain YFP, the synthetized SH2 domain was inserted into pCri-7b and 8b (previously linearized with XhoI) by Gibson assembly to construct EZ-L664 and EZ-L703 (see Supplementary table 4). Monobody HA4 was inserted into pCri-7b; the vector was digested (opened) with NheI and XhoI and Gibson-assembled to build the template (EZ-L663) used for the AsLOV2 insertions. A stop codon was added before the in-frame (C-terminus) 6x-histidine tag of pCri-7b. The AsLOV2 domain (residues 408-543) was either amplified by PCR from previous constructs^45^ (wt AsLOV2), or synthesized as IDT gBlocks (AsLOV2 mutants). To insert AsLOV2 into the monobody, the backbone from the initial construct containing HA4 (EZ-L663) was PCR amplified, using Takara Hifi PCR premix, starting from the insertion positions (Supplementary Table 1) that were selected (and adding homology arms to AsLOV2). Next, chimeras were finally assembled mixing each of the amplified products of the backbone PCR from EZ-L663 with the AsLOV2 domain obtained from either PCR amplification (wt AsLOV2) or synthesized by gBlocks (AsLOV2 mutants). SUMO tags were added by inserting (with Gibson assembly) the PCR product of the full-length chimeras (with homology arms) into the pCri-11b plasmid, previously opened with NheI and XhoI. *PDC1* was amplified from *S. cerevisiae* (S288C) genomic DNA using PCR, also with homology arms, and the construct (EZ-L886) was built via Gibson assembly (3 fragments) with digested pCri-7b (NheI and XhoI) and the PCR product of *SH2* amplified from EZ-L664 (with homology arms). All constructs (Supplementary Table 4) were sequenced by Genewiz and all protein sequences are available in Supplementary Sequences. We used chemically competent DH5α to clone all vectors. After verifying the plasmid sequence, vectors were used to transform chemically competent BL21 (DE3) or Rosetta strains for protein expression.

### Construction of the structural model of OptoMB

To build the structural model of the HA4-AsLOV2 chimera (OptoMB) interacting with the SH2 domain (Fig. 1e), a shortened version^45^ (residues 408-543) of the AsLOV2 domain (PDB ID: 2V1A)^64^ was manually inserted to residues S58 and S59 of the monobody HA4, using the program *Coot*^65^. The crystal structure of HA4 in complex with the SH2 domain (PDB ID: 3k2m)^46^ was used as template. After a manual adjustment, an energy minimization of the HA4-AsLOV2 chimera was carried out with the website version of *YASARA*^66^ (http://www.yasara.org/minimizationserver.htm).

### Plasmid Construction for mammalian cells

Constructs for mammalian cell experiments were cloned using backbone PCR and inFusion (Clontech). Monobody HA4 or OptoMB variants were PCR amplified and Gibson-assembled from bacterial plasmids (described above) into a pHR vector with a C-terminal irFP fusion (Addgene #111510). The SH2 domain was amplified from EZ-L664 using PCR, and Gibson-assembled into a pHR vector containing a C-terminal mCherry-CAAX fusion tag (Addgene #50839). Stellar *E. coli* cells (TaKaRa) were transformed with these plasmids for amplification and DNA storage. All plasmids were sequenced by Genewiz to verify quality.

### Lentivirus Production and Transduction

HEK 293T cells were plated on a 12-well plate, reaching 40% confluency the next day. The cells were then co-transfected with the corresponding pHR plasmid and lentiviral packaging plasmids (pMD and CMV) using Fugene HD (Promega). Cells were incubated for ~ 48 hours and virus was collected and filtered through a 0.45 mm filter. In addition, 2 μL of polybrene and 40 μL of HEPES were added to the 2 mL viral solution. For infection, HEK 293T cells were plated on a 6-well plate, allowed to adhere and reach 40% confluency. At that time, the cells were infected with 200-500 μL of viral solution. All imaging was done at least 48 hours post infection. Cells were cultured in DMEM media with 10% FBS, penicillin (100 U/mL), and streptomycin (0.1 mg/mL).

### Screening for light-responsive monobodies

A 6x-histidine tagged fusion of YFP and SH2 (His_6_-YFP-SH2) was grown in 500 mL of autoinduction^67^ media + Kanamycin (Kan)(50 μg/mL) for 16 hours at 30°C. Monobody HA4 and monobody-AsLOV2 chimeras were grown in 250 mL of autoinduction media + Kan for 16 hours at 30°C. For each test, monobody HA4 (used as control) and 3 different monobody-AsLOV2 chimeras were tested simultaneously. Cells were then harvested by centrifugation at 7500 x*g* for 20 minutes at 4°C in a Lynx 4000 centrifuge (Sorvall™) and supernatant was discarded. Cell pellets were resuspended in Binding buffer (Tris 100 mM pH 8.0, NaCl 150 mM, Glycerol 1% and 5 mM imidazole) supplemented with 1 mM of PMSF, adding 8 mL to the cells containing His_6_-YFP-SH2 and 3 mL for the monobody HA4 and each of the chimeras. The resuspended cells were flash-frozen forming droplets directly into liquid nitrogen (LN2) and placed in small (LN2-cold) grinding vials to be disrupted using a CryoMill system (Spex sample prep®) with a cycle of 2 min grinding and 3 min cooling (14 times). The broken cell powder was thawed in 50 mL Falcon tubes at room temperature with the addition of 4 mL of Lysis buffer (Binding buffer supplemented with 1 mM PMSF and 2 mg/mL of DNAse) for His_6_-YFP-SH2 and 2 mL of Lysis buffer for monobody HA4 and each of the chimeras. Once thawed, the bacterial lysates were centrifuged at 25,000 ×*g* in a Lynx 4000 centrifuge (Sorvall™) for 30 minutes at 4°C and the supernatant (clarified lysate) was transferred to 15 mL conical tubes. To enhance AsLOV2 activity, we added flavin mononucleotide (FMN) to a final concentration of 0.25 mg/ml, to each of the chimera-containing lysate and then incubated for 15 minutes at 4°C by vertical rotation in a tube Revolver/Rotator at 50 rpm (Thermo Scientific™). The His_6_-YFP-SH2 supernatant was then mixed with 4.5 mL of a 50% suspension of Co-charged agarose resin (Talon^®^), previously equilibrated in Binding buffer, and incubated at 4°C, rocking for 45 min. The suspension was sedimented by gravity and the supernatant discarded. The beads were then thoroughly washed with Binding buffer. with approximately 50 times the resin or column volume. The beads (now with His_6_-YFP-SH2 bound) were then finally resuspended in Binding buffer to a total volume of 13.5 mL. The bead suspension was equally divided into 9 conical tubes of 15 mL (having each 1.5 mL of the bead suspension). Four of these tubes were used for experiments under blue light with LED panels (450 nM) while the other four were used for experiments in the dark (wrapped in aluminum foil). Experiments were performed in a dark room and red light was used occasionally for visualization purposes. The remaining tube was left as a control for His_6_-YFP-SH2 binding. Samples of 2.5 mL of bacterial clarified lysates containing either the monobody HA4 supernatant or each one of the chimeras were added to separate tubes of resin. The mixtures were then incubated for 45 minutes (at 4°C and constant vertical rotation of 20 rpm) under blue light or dark conditions. Tubes were then allowed to settle at 4°C under the same light conditions in which they were incubated, for 15-20 minutes. After carefully discarding the supernatants, the beads were washed 5 to 6 times; each time with 10 mL of Binding buffer and rotating at 20 rpm for 15 min. After each wash, the mixtures were allowed to settle at 4°C for 15-20 minutes under the same light conditions as they were incubated and the supernatants were again discarded. This step was repeated until beads were washed with approximately 40-50 resin-volumes. During the course of the experiments, light conditions within the room were carefully held constant and red light was for visualization purposes used only when needed, specially to minimize any blue light exposure of the dark samples when opening the wrapped tubes to exchange buffer (washing, incubation and elution). After the last wash, proteins were eluted with 2.5 mL of Elution buffer (Tris 100 mM pH 8.0, 150 mM NaCl, 1% Glycerol, 500 mM Imidazole) and then equal volumes of elution samples (dark and light) for each chimera (and control) were loaded onto 12% polyacrylamide gels and resolved with SDS-PAGE.

### Purification of Monobody HA4, OptoMB (and all its variants), and YFP-SH2

HA4, monobody-AsLOV2 chimeras (OptoMB and variants), and YFP-SH2 constructs were purified using N-terminal 6x-histidine tag fusions and metal affinity chromatography. Chimeras were protected from excessive ambient light exposure to prevent potential protein destabilization and improve yields. This included covering the shakers with black blankets during expression, wrapping culture flasks and tubes (containing crude or purified proteins) with thick aluminum foil and performing the chromatography in the dark or with red light (when needed). OptoMB (and all its variants) were expressed at 18°C for three days in 1 L or 2 L of Autoinduction media^67^ (plus Kan 50 μg/mL). HA4 and YFP-SH2 were expressed in 1 L or 2 L of Autoinduction media (plus Kan 50 μg/mL) for 16 h at 30°C. Cells were then harvested by centrifugation at 7500 ×*g* for 20 minutes at 4°C in a Lynx 6000 centrifuge (Sorvall™) and supernatant was discarded. Each cell pellet was resuspended in between 8 to 12 mL of Binding buffer and frozen droplets were prepared as described above and immediately transferred to a large (LN2-cold) grinding tube. Cryogenic grinding was performed as described above. Broken cells (frozen powder) were thawed in 50 mL Falcon tubes at room temperature with the addition of Lysis buffer up to 5% of the initial cell culture volume. After clarifying the bacterial lysates by centrifugation as described above, these were loaded onto columns of 2 to 5 mL (50% suspension) of Co-charged resin (Talon^®^) previously equilibrated with Binding buffer (Tris 100 mM pH 8.0, NaCl 150 mM, Glycerol 1% and 5 mM Imidazole). Columns were washed with 40-50 column volumes of Binding buffer and proteins eluted with Elution buffer (Binding buffer supplemented with 250 mM of imidazole). Proteins were then run through size exclusion chromatography (SEC) using a Hiprep™ HR 16/60 Sephacryl™ 200 with Buffer A (Tris 50 mM pH 8.0 and NaCl 150 mM), in an FPLC (ÄKTA pure from GE^®^ Healthcare). The aliquots enriched with the target proteins were concentrated by centrifugation (in cycles of 10 min) at maximum speed in a Sorvall Legend XTR Benchtop centrifuge (Thermo Scientific™) using 15 mL centricons^®^ (Millipore) with a cutoff selected according to the molecular size of each protein, until concentrations between 3 to 8 mg/mL were reached.

### Size-exclusion chromatography to characterize OptoMB-SH2 interaction in solution

Purified monobody HA4 (75 μL from a 202 μM sample) or OptoMB (wt AsLOV2) (300 μL from a 40.7 μM sample) dissolved in Buffer A, were mixed with YFP-SH2 (200 μL from a 58.8 μM sample in Buffer A) in approximately 1.2:1 molar ratio of binder to target. Each mixture was incubated for 10 min in either dark or blue light (450 nm) before loading the full volume (275 or 500 μl) onto a Superdex™ 200 16/300 column (GE^®^ Healthcare), which was then run at 1 mL/min with Buffer A at 4°C (using an ÄKTA pure from GE^®^ Healthcare). To test different light conditions, the column was either illuminated with wrapped blue LED strips (450 nm) or covered with aluminum foil for the total duration of the filtration (Supplementary Fig. 2c). For experiments in the dark, we also minimized light sources in the room and covered the chromatography cabinet with a black blanket. The SEC was monitored by UV absorbance at 280 nm.

### Intracellular imaging of HEK293T cells

For live cell imaging, we used 0.17 mm glass-bottomed black walled 96-well plates (In Vitro Scientific). Glass was first treated with 10 μg/mL of fibronectin in PBS for 20 min. HEK293T cells expressing both SH2-mCherry-CAAX and either the monobody HA4 or OptoMB were then plated and allowed to adhere onto the plate before imaging. Mineral oil (50 μL) was added on top of each well with cells prior to imaging, to limit media evaporation. The mammalian cells were kept at 37°C with 5% CO_2_ for the duration of the imaging experiments. The irFP and mCherry fluorescence were imaged using a Nikon Eclipse Ti microscope with a Prior linear motorized stage, a Yokogawa CSU-X1 spinning disk, an Agilent laser line module containing 405, 488, 561 and 650 nm lasers, an iXon DU897 EMCCD camera, and a 40X oil immersion objective lens. An LED light source was used for photoexcitation with blue light (450 nm), which was delivered through a Polygon400 digital micro-mirror device (DMD; Mightex Systems).

### Imaging of coated Agarose Beads

Approximately 200 μL of Ni-NTA agarose slurry (50% suspension) (Qiagen) equilibrated in Buffer A were mixed with 500 μL of 100 μM of either monobody HA4 or OptoMB (AsLOV2 V416L variant) in an Eppendorf tube (covered with aluminum foil) and incubated by vertical rotation (at 20 rpm for 20 min) at room temperature to allow binding through the 6x-histidine tag until saturation. The excess protein was then washed twice with 1 mL of Buffer A, centrifuging the beads at low speed for 1 min (1000 rpm in a benchtop centrifuge) each time; finally discarding the excess of supernatant after the second wash. Then, 50 μL of a purified YFP-SH2 solution at 2 μM (with the 6x-histidine tag cleaved off) was added onto 0.17 mm glass-bottomed black walled 96-well plate (In Vitro Scientific) followed by 2 μL of the washed resin with the beads labeled with OptoMonobody (or monobody HA4 as control). The mixture was equilibrated for at least 1 hour at room temperature and up to overnight at 4°C prior to imaging, performed at room temperature. The same microscope setup as described for the cells imaging was used, except for the objective (20X in this case), to follow YFP fluorescence over time on the surface of the bead. For the spatial control of the OptoMB-SH2 interaction on beads the same setup was used. Two beads in an area of around 200 × 250 μm were imaged at the same time applying a light (450 nm) mask, which uses a circular ROI with a radius of 70 μm to cover and illuminate only one bead. The YFP fluorescence was recorded over time (for a total of 1h) for both, the illuminated and the unilluminated bead. Quantification was performed by measuring the change in YFP fluorescence intensity over time in a defined region on the surface of the bead (using ImageJ^68^) and subtracting the background.

### Calculation of binding kinetics by Bio-layer interferometry

Measurements of the binding (*k*_on_) and unbinding (*k*_off_) rate constants, as well as the dissociation (or affinity) constant (*K*_d_) for HA4 monobody and OptoMB (including variants V416L and V416I-G528A-N538E) were performed on Octet RED96e instruments (ForteBio). Ni-NTA sensors (ForteBio) were first equilibrated using Buffer A for 10 min prior the measurement. A volume of 200 μL of Buffer A or protein solutions (previously dialyzed with Buffer A when needed) were added to clear 96-well plates. During the experimental run, the Ni-NTA sensors were first immersed in Buffer A to record the baseline. Protein binders were then loaded by switching to wells with solutions of 6x-histidine tagged HA4 monobody or OptoMB variants (with concentrations between 100 μg/mL and up to 1 mg/mL) until values of ~ 4 nm were reached (avoiding saturation of the sensors). The sensors were then transferred back into Buffer A to remove unbound protein. To measure the binding rate constant (*k*_on_) the sensors with bound monobody HA4 or OptoMB variants were subsequently shifted to wells containing various concentrations of YFP-SH2 (at concentrations indicated Fig 2d-e and Supplementary Fig. 2e-f). To measure the unbinding rate constant (*k*_off_), the sensors were then moved to wells containing Buffer A to trigger dissociation of YFP-SH2. To measure binding kinetics of the light state, the lid of the Octet remained open during the measurement and a blue LED panel was held above the 96-well plate, maintaining constant blue illumination for the duration of the experiment. For the OptoMB variant with the AsLOV2 mutations V416I-G528A-N538E, lit states were sufficiently long-lived to remain fully activated in response to pre-illumination with a blue light panel and the continuous illumination of the Octet sensors. The raw binding and unbinding data were simultaneously fit to models of the binding and unbinding reactions:

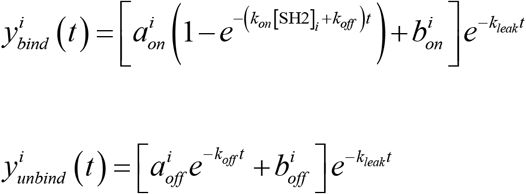

This model incorporates the following dependent and independent variables:

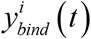 refers to the i^th^ binding curve
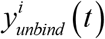 the i^th^ unbinding curve
[SH2]_*i*_ refers to the concentration of YFP-SH2 used for the i^th^ binding curve
*t* is the time elapsed since the start of the binding/unbinding phase.

It also includes the following parameters:

*k_on_* is the on-rate (enforced to be identical across all binding and unbinding curves)
*k_off_* is the off-rate (enforced to be identical across all binding and unbinding curves)
*k_leak_* represents the slow unbinding of His-tagged OptoMB from the probe, leading to a gradual decay of signal throughout the entire experiment.
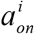 is the total change in signal due to SH2 binding for the i^th^ curve
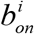 is the signal baseline during the binding phase for the i^th^ curve
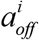 is the total change in signal due to SH2 unbinding for the i^th^ curve
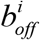 is the signal baseline during the unbinding phase for the i^th^ curve

The model thus contains 4\**n* + 3 parameters, where *n* is the number of distinct SH2 concentrations tested. Fits were performed using nonlinear gradient descent using the MATLAB `fmincon` function. Code is available as described in Ref. 45.

### Light-controlled affinity chromatography (LCAC) of SH2-tagged proteins using cobalt-immobilized OptoMB

To prepare our OptoMB resin and columns, we incubated the purified variant OptoMB_V416I_G528A_N538E_ (having a SUMO tag in the N-term to boost expression, Supplementary Table 4) with Co-charged resin (Talon^®^) (50% suspension) previously equilibrated in Buffer A, in order to produce 2.5 mL of resin with approximately 8 mg of OptoMB bound per mL. The Resin was rotated in 50 mL conical tubes (covered with aluminum foil) at room temperature for 20 min and then poured in 20 mL BioRad glass Econo-Columns and washed with Buffer A (50 column volumes). The use of imidazole in the Buffer A was avoided because it interferes with the photocycle of the AsLOV2 domain, enhancing the rate of recovery of the dark state thus reducing the lifetime of the lit state^69^, which would reduce the efficiency of light elution. Production of the OptoMB resin (and columns) and the LCAC procedure were carried out at room temperature within a carefully controlled dark room to avoid blue light contamination, using red light from LED panels occasionally for visualization purposes. Approximately 10 to 12 mL of clarified bacterial lysates (obtained as described before from 1L cultures) containing YFP-SH2 or SH2-PDC1 (without his-tags) were incubated with 2.5 mL of OptoMB resin in Buffer A, which was enough to saturate the OptoMB resin with binding targets. The mixture was then vertically rotated gently for 45 min in 15 ml conical tubes. The OptoMB resin was then poured into 10 mL BioRad glass Econo-Columns or left to settle to discard the supernatant by pipetting (experiments in batch). Approximately 80 column volumes of Buffer A were used to wash the OptoMB resin. When purifying in batch, 40 mL of Buffer A were added to the resin at a time in 50 mL conical tubes; after rotating for 5 min, the resin was left to settle to remove the supernatant by pipetting. This cycle was repeated until reaching 80 resin volumes. After washing, the resin was poured back into a 15 mL conical tube. When purifying in batch, for each light-elution step we added 2-times the resin volume of Buffer A and incubated with gentle rocking in front of a blue light (450 nm) LED panel (approximately 20 cm away) for 10 to 15 min; then letting the resin settle while illuminated and recovering the light-eluted aliquot by pipetting. When purifying in columns, 2 column bed volumes of Buffer A were poured at a time while the column was exposed to blue light from three LED panels in a triangular conformation (surrounding the column), and collected by gravity. For both batch and column purifications, the elution steps were repeated until no more protein eluted from the resin (usually 4 to 5 times). In both methods, batch or column, after light-elution we added 4 times the resin volume of Buffer A containing 250 mM of imidazole to elute the proteins still bound to the resin after light-elution, needed to calculate yields and resin capacity. Samples from the different purification steps were resolved with SDS-PAGE, using 12% SDS-polyacrylamide gels.

### Light-controlled affinity chromatography (LCAC) of SH2-tagged proteins using covalently coupled OptoMB

A total of 1-1.5g (dry weight) of CNBr-activated Sepharose™ 4B (GE^®^ Healthcare) was resuspended in Conjugation buffer (NaCHO_3_ 0.1M pH 8.3 and 500 mM NaCl) according to the manufacturer specifications. The excess of Conjugation buffer was discarded and the hydrated resin mixed with purified OptoMB_V416I_G528A_N538E_ (containing an N-terminal SUMO tag to boost expression, Supplementary Table 4), previously dialyzed against Conjugation buffer by centrifugation (using 15 mL centricons^®^ as describe above). For each preparation, 5 mL (50% suspension) of resin was conjugated with OptoMB at approximately 8 mg of OptoMB per mL of resin. The manufacturer’s protocol was followed with minor modifications, keeping all protein immobilization steps (conjugation, blockage and acid/base washing cycles) in the dark by covering the tubes with thick aluminum foil. The acid/base washing cycles were reduced to one cycle, as the three washes recommended by the manufacturer results in loss of the FMN, the AsLOV2 co-factor (as evidenced by the loss of the protein’s yellow coloration immediately after the second wash). Once OptoMB was conjugated, the resin was washed and equilibrated with Buffer AS (Buffer A with salt concentration increased to 300 mM NaCl). The OptoMB-conjugated resins were used immediately or stored at 4°C (protected from light with foil), where they are stable for up to at least a week (yielding same results as fresh resin). To purify YFP-SH2 or SH2-PDC1, approximately 10 to 15 mL of the clarified bacterial lysates obtained from 1 L cultures (as described above) were incubated in batch with 5 mL (50% suspension) of the OptoMB-conjugated sepharose resin, which was enough to saturate the OptoMB resin with binding targets. The same protocol for LCAC purification with OptoMB-immobilized in Co-charged (Talon^®^) in batch described above was followed with CNBr-OptoMB resin, except Buffer AS was used instead of Buffer A. To calculate the yields and resin capacity, after the final light-elution step, 200 μl of the 50% resin suspension were resuspended in 100 μl of SDS-PAGE sample-loading buffer and incubated at 100°C for 10 min in a heat block, before loading onto the gel. Samples from all purification steps were resolved on a 12% SDS-polyacrylamide gels.

### Calculations of Protein Purity, Yield and Resin capacity

Protein purity was estimated using densitometric analysis in the FIJI implementation of ImageJ^68^. The software analyzes individual bands from a scanned gel, returning the integrated intensity of each band. For each aliquot of purified protein (eluted with light), we selected the band representing the SH2-tagged protein, and used their integrated intensities to calculate the fraction of the purified target versus the overall total of protein present in that fraction.

The overall yield (*y*) represents the total amount (mg) of purified protein that eluted with light (*TPE*) as a percentage of the total amount (mg) of protein that was initially bound to the resin (*TPB*). *TPE* was calculated by multiplying the concentration (mg/mL) of all the light-eluted fractions ([*TPE*]) by their total volume (mL) (*TPEvol*). The concentrations of the fractions eluted with light were calculated using Absorbance at 280 nm with a spectrometer (Biospectrometer, Eppendorf). *TPB* was equal to *TPE* plus the total residual SH2-tagged protein remaining on the column after light-elution (*TPR*). To calculate the *TPR*, we recovered all residual protein using imidazole elution (of Co-Agarose-OptoMB) or heat treatment (of CNBr-OptoMB) of the beads after light-elution. Concentrations of recovered residual protein were calculated by running each sample on a 12% polyacrylamide gel alongside standards of the same SH2-tagged protein of known concentrations. Band intensities were then computed by densitometry analysis in FIJI as above. The amount of target protein still bound to the column was thus equal to the concentration retrieved in this final elution [*TRP*] multiplied by its total elution volume *TRPvol*. Finally, the resin capacity *C_R_* was calculated as *TPB* divided by the total volume of the resin used *V_R_*.

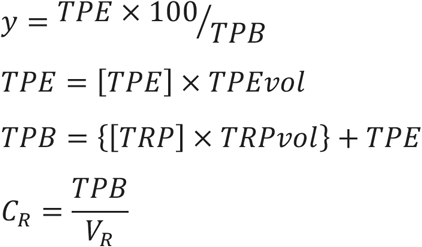

Where

*y* is the yield of purified protein obtained with the resin
*TPE* is the total amount of protein eluted with light
[*TPE*] is the concentration of all the fractions that eluted with light
*TPEvol* is the total volume of the light-elution fraction
*TPB* is the total amount of protein initially bound to the resin
[*TRP*] is the concentration of the total residual protein fraction that was still bound to the resin after the elution with light
*TRPvol* is the total volume of the fraction containing the total residual protein that was still bound to the resin after the elution with light, and was recovered by imidazole or heat treatment of the beads.
*C_R_* is the protein binding capacity of the resin
*V_R_* is the volume of resin used in the experiment

## Data Availability

The authors declare that all data supporting the findings of this study are available within the paper (and its supplementary information files), but original data that supports the findings are available from the corresponding authors upon reasonable request.

## Author contributions

Conceptualization, C.C.L., E.M.Z., A.A.G., J.E.T. and J.L.A.; Methodology, C.C.L., E.M.Z., A.A.G., J.E.T. and J.L.A.; Investigation, C.C.L., E.M.Z, A.A.G, and N.A.; Writing – Original Draft, C.C.L., E.M.Z. and J.L.A.; Review & Editing, all authors; Funding Acquisition, J.E.T. and J.L.A.; Supervision, J.E.T. and J.L.A.

## Acknowledgements

We thank all members of Avalos and Toettcher labs for helpful comments. We also thank the Biophysics Core Facility and especially Venu Vandavasi for help with the bio-layer interferometry measurements. J.E.T. was supported by NIH grant DP2EB024247 and J.L.A was supported by the U.S. Department of Energy, Office of Science, Office of Biological and Environmental Research Award Number DE-SC0019363, the NSF CAREER Award CBET-1751840, The Pew Charitable Trusts, The Eric and Wendy Schmidt Transformative Technology Fund Award, and the Camille Dreyfus Teacher-Scholar Award.

