## Supplementary Information for "Light-responsive monobodies for dynamic control of customizable protein binding"

**Supplementary Sequences:**

The AsLOV2 sequence is highlighted in blue.

HA4-AsLOV2 (SS58 insertion):
GSSVSSVPTKLEVVAATPTSLLISWDAPMSSSSVYYYRITYGETGGNSPVQEFTVPYSGLERIEKNFVITDPRLPDNPIIFASDSFLQLTEYSREEILGRNCRFLQGPETDRATVRKIRDAIDNQTEVTVQLINYTKSGKKFWNLFHLQPMRDQKGDVQYFIGVQLDGTEHVRDAAEREGVMLIKKTAENIDEAAGSSTATISGLSPGVDYTITVYAWGEDSAGYMFMYSPISINYRTC*

HA4-AsLOV2 (MS29 insertion with residues removed from L1):
GSSVSSVPTKLEVVAATPTSLLISWDAPMGLERIEKNFVITDPRLPDNPIIFASDSFLQLTEYSREEILGRNCRFLQGPETDRATVRKIRDAIDNQTEVTVQLINYTKSGKKFWNLFHLQPMRDQKGDVQYFIGVQLDGTEHVRDAAEREGVMLIKKTAENIDEAAGSVYYYRITYGETGGNSPVQEFTVPYSSSTATISGLSPGVDYTITVYAWGEDSAGYMFMYSPISINYRTC*

HA4-AsLOV2 (MS29 insertion):
GSSVSSVPTKLEVVAATPTSLLISWDAPMGLERIEKNFVITDPRLPDNPIIFASDSFLQLTEYSREEILGRNCRFLQGPETDRATVRKIRDAIDNQTEVTVQLINYTKSGKKFWNLFHLQPMRDQKGDVQYFIGVQLDGTEHVRDAAEREGVMLIKKTAENIDEAAGSSSSVYYYRITYGETGGNSPVQEFTVPYSSSTATISGLSPGVDYTITVYAWGEDSAGYMFMYSPISINYRTC*

HA4-AsLOV2 (GN46 insertion with residues removed from L3):
GSSVSSVPTKLEVVAATPTSLLISWDAPMSSSSVYYYRITYGETGGLERIEKNFVITDPRLPDNPIIFASDSFLQLTEYSREEILGRNCRFLQGPETDRATVRKIRDAIDNQTEVTVQLINYTKSGKKFWNLFHLQPMRDQKGDVQYFIGVQLDGTEHVRDAAEREGVMLIKKTAENIDEAAGNSPVQEFTVPYSSSTATISGLSPGVDYTITVYAWGEDSAGYMFMYSPISINYRTC*

HA4-AsLOV2 (TN45 insertion with residues removed from L3):
GSSVSSVPTKLEVVAATPTSLLISWDAPMSSSSVYYYRITYGETGLERIEKNFVITDPRLPDNPIIFASDSFLQLTEYSREEILGRNCRFLQGPETDRATVRKIRDAIDNQTEVTVQLINYTKSGKKFWNLFHLQPMRDQKGDVQYFIGVQLDGTEHVRDAAEREGVMLIKKTAENIDEAAGNSPVQEFTVPYSSSTATISGLSPGVDYTITVYAWGEDSAGYMFMYSPISINYRTC*

HA4-AsLOV2 (SP68 insertion with residues removed from L5):
GSSVSSVPTKLEVVAATPTSLLISWDAPMSSSSVYYYRITYGETGGNSPVQEFTVPYSSSTATISGSGLERIEKNFVITDPRLPDNPIIFASDSFLQLTEYSREEILGRNCRFLQGPETDRATVRKIRDAIDNQTEVTVQLINYTKSGKKFWNLFHLQPMRDQKGDVQYFIGVQLDGTEHVRDAAEREGVMLIKKTAENIDEAAGPVDYTITVYAWGEDSAGYMFMYSPISINYRTC*

HA4-AsLOV2 (PT18 insertion with residues removed from L1):
GSSVSSVPTKLEVVAAPGLERIEKNFVITDPRLPDNPIIFASDSFLQLTEYSREEILGRNCRFLQGPETDRATVRKIRDAIDNQTEVTVQLINYTKSGKKFWNLFHLQPMRDQKGDVQYFIG

VQLDGTEHVRDAAEREGVMLIKKTAENIDEAAGTSLLISWDAPMSSSSVYYYRITYGETGGNSPVQEFTVPYSSSTATISGLSPGVDYTITVYAWGEDSAGYMFMYSPISINYRTC*

AsLOV2 domain:
GLERIEKNFVITDPRLPDNPIIFASDSFLQLTEYSREEILGRNCRFLQGPETDRATVRKIRDAIDNQTEVTVQLINYTKSGKKFWNLFHLQPMRDQKGDVQYFIGVQLDGTEHVRDAAEREGVMLIKKTAENIDEAAG

Monobody HA4:
GSSVSSVPTKLEVVAATPTSLLISWDAPMSSSSVYYYRITYGETGGNSPVQEFTVPYSSSTATISGLSPGVDYTITVYAWGEDSAGYMFMYSPISINYRTC*

$His_6$-YFP-SH2:
HHHHHHSSGENLYFQGHMASKVSKGEELFTGVVPILVELDGDVNGHKFSVSGEGEGDATYGKLTLKFICTTGKLPVPWPTLVTTFGYGLQCFARYPDHMKQHDFFKSAMPEGYVQERTIFFKDDGNYKTRAEVKFEGDTLVNRIELKGIDFKEDGNILGHKLEYNYNSHNVYIMADKQKNGIKVNFKIRHNIEDGSVQLADHYQQNTPIGDGPVLLPDNHYLSTQSALSKDPNEKRDHMVLLEFVTAAGITLGMDELYKSLEKHSWYHGPVSRNAAEYLLSSGINGSFLVRESESSPGQRSISLRYEGRVYHYRINTASDGKLYVSSESRFNTLAELVHHHSTVADGLITTLHYPAPKRNKPTVYGVSPNY*

HA4-AsLOV2 (SS30 insertion):
GSSVSSVPTKLEVVAATPTSLLISWDAPMSGLERIEKNFVITDPRLPDNPIIFASDSFLQLTEYSREEILGRNCRFLQGPETDRATVRKIRDAIDNQTEVTVQLINYTKSGKKFWNLFHLQPMRDQKGDVQYFIGVQLDGTEHVRDAAEREGVMLIKKTAENIDEAAGSSSVYYYRITYGETGGNSPVQEFTVPYSSSTATISGLSPGVDYTITVYAWGEDSAGYMFMYSPISINYRTC*

HA4-AsLOV2 (NS47 insertion):
GSSVSSVPTKLEVVAATPTSLLISWDAPMSSSSVYYYRITYGETGGNGLERIEKNFVITDPRLPDNPIIFASDSFLQLTEYSREEILGRNCRFLQGPETDRATVRKIRDAIDNQTEVTVQLINYTKSGKKFWNLFHLQPMRDQKGDVQYFIGVQLDGTEHVRDAAEREGVMLIKKTAENIDEAAGSPVQEFTVPYSSSTATISGLSPGVDYTITVYAWGEDSAGYMFMYSPISINYRTC*

HA4-AsLOV2 (SA84 insertion):
GSSVSSVPTKLEVVAATPTSLLISWDAPMSSSSVYYYRITYGETGGNSPVQEFTVPYSSSTATISGLSPGVDYTITVYAWGEDSGLERIEKNFVITDPRLPDNPIIFASDSFLQLTEYSREEILGRNCRFLQGPETDRATVRKIRDAIDNQTEVTVQLINYTKSGKKFWNLFHLQPMRDQKGDVQYFIGVQLDGTEHVRDAAEREGVMLIKKTAENIDEAAGAGYMFMYSPISINYRTC*

YFP-SH2:
MASKVSKGEELFTGVVPILVELDGDVNGHKFSVSGEGEGDATYGKLTLKFICTTGKLPVPWPTLVTTFGYGLQCFARYPDHMKQHDFFKSAMPEGYVQERTIFFKDDGNYKTRAEVKFEGDTLVNRIELKGIDFKEDGNILGHKLEYNYNSHNVYIMADKQKNGIKVNFKIRHNIEDGSVQLADHYQQNTPIGDGPVLLPDNHYLSTQSALSKDPNEKRDHMVLLEFVTAAGITLGMDELYKSLEKHSWYHGPVSRNAAEYLLSSGINGSFLVRESESSPGQRSISLRYEGRVYH

YRINTASDGKLYVSSESRFNTLAELVHHHSTVADGLITTLHYPAPKRNKPTVYGVSPNY*

His$_6$-HA4:
HHHHHHSSGENLYFQGHASGSSVSSVPTKLEVVAATPTSLLISWDAPMSSSSVYYYRITY
GETGGNSPVQEFTVPYSSSTATISGLSPGVDYTITVYAWGEDSAGYMFMYSPISINYRTC
*

HA4-AsLOV2 (TG44 insertion):
GSSVSSVPTKLEVVAATPTSLLISWDAPMSSSSVYYYRITYGETGLERIEKNFVITDPRLP
DNPIIFASDSFLQLTEYSREEILGRNCRFLQGPETDRATVRKIRDAIDNQTEVTVQLINYTK
SGKKFWNLFHLQPMRDQKGDVQYFIGVQLDGTEHVRDAAEREGVMLIKKTAENIDEA
AGGGNSPVQEFTVPYSSSTATISGLSPGVDYTITVYAWGEDSAGYMFMYSPISINYRTC*

HA4-AsLOV2 (SG65 insertion):
GSSVSSVPTKLEVVAATPTSLLISWDAPMSSSSVYYYRITYGETGGNSPVQEFTVPYSSST
ATISGLERIEKNFVITDPRLPDNPIIFASDSFLQLTEYSREEILGRNCRFLQGPETDRATVRK
IRDAIDNQTEVTVQLINYTKSGKKFWNLFHLQPMRDQKGDVQYFIGVQLDGTEHVRDA
AEREGVMLIKKTAENIDEAAGGLSPGVDYTITVYAWGEDSAGYMFMYSPISINYRTC*

HA4-AsLOV2 (DA26 insertion):
GSSVSSVPTKLEVVAATPTSLLISWDGLERIEKNFVITDPRLPDNPIIFASDSFLQLTEYSRE
EILGRNCRFLQGPETDRATVRKIRDAIDNQTEVTVQLINYTKSGKKFWNLFHLQPMRDQ
KGDVQYFIGVQLDGTEHVRDAAEREGVMLIKKTAENIDEAAGAPMSSSSVYYYRITYGE
TGGNSPVQEFTVPYSSSTATISGLSPGVDYTITVYAWGEDSAGYMFMYSPISINYRTC*

HA4-AsLOV2 (ED82 insertion):
GSSVSSVPTKLEVVAATPTSLLISWDAPMSSSSVYYYRITYGETGGNSPVQEFTVPYSSST
ATISGLSPGVDYTITVYAWGEGLERIEKNFVITDPRLPDNPIIFASDSFLQLTEYSREEILGR
NCRFLQGPETDRATVRKIRDAIDNQTEVTVQLINYTKSGKKFWNLFHLQPMRDQKGDV
QYFIGVQLDGTEHVRDAAEREGVMLIKKTAENIDEAAGDSAGYMFMYSPISINYRTC*

HA4-AsLOV2 (DS83 insertion):
GSSVSSVPTKLEVVAATPTSLLISWDAPMSSSSVYYYRITYGETGGNSPVQEFTVPYSSST
ATISGLSPGVDYTITVYAWGEDGLERIEKNFVITDPRLPDNPIIFASDSFLQLTEYSREEILG
RNCRFLQGPETDRATVRKIRDAIDNQTEVTVQLINYTKSGKKFWNLFHLQPMRDQKGD
VQYFIGVQLDGTEHVRDAAEREGVMLIKKTAENIDEAAGSAGYMFMYSPISINYRTC*

HA4-AsLOV2 (GG45 insertion):
GSSVSSVPTKLEVVAATPTSLLISWDAPMSSSSVYYYRITYGETGGLERIEKNFVITDPRL
PDNPIIFASDSFLQLTEYSREEILGRNCRFLQGPETDRATVRKIRDAIDNQTEVTVQLINYT
KSGKKFWNLFHLQPMRDQKGDVQYFIGVQLDGTEHVRDAAEREGVMLIKKTAENIDE
AAGGNSPVQEFTVPYSSSTATISGLSPGVDYTITVYAWGEDSAGYMFMYSPISINYRTC*

HA4-AsLOV2 (PT18 insertion):
GSSVSSVPTKLEVVAATPGLERIEKNFVITDPRLPDNPIIFASDSFLQLTEYSREEILGRNCR
FLQGPETDRATVRKIRDAIDNQTEVTVQLINYTKSGKKFWNLFHLQPMRDQKGDVQYFI

GVQLDGTEHVRDAAEREGVMLIKKTAENIDEAAGTSLLISWDAPMSSSSVYYYRITYGETGGNSPVQEFTVPYSSSTATISGLSPGVDYTITVYAWGEDSAGYMFMYSPISINYRTC*

HA4-AsLOV2 (SP68 insertion):
GSSVSSVPTKLEVVAATPTSLLISWDAPMSSSSVYYYRITYGETGGNSPVQEFTVPYSSSTATISGLSGLERIEKNFVITDPRLPDNPIIFASDSFLQLTEYSREEILGRNCRFLQGPETDRATVRKIRDAIDNQTEVTVQLINYTKSGKKFWNLFHLQPMRDQKGDVQYFIGVQLDGTEHVRDAAEREGVMLIKKTAENIDEAAGPGVDYTITVYAWGEDSAGYMFMYSPISINYRTC*

AsLOV2-HA4 (N-terminus):
GLERIEKNFVITDPRLPDNPIIFASDSFLQLTEYSREEILGRNCRFLQGPETDRATVRKIRDAIDNQTEVTVQLINYTKSGKKFWNLFHLQPMRDQKGDVQYFIGVQLDGTEHVRDAAEREGVMLIKKTAENIDEAAGGSSVSSVPTKLEVVAATPTSLLISWDAPMSSSSVYYYRITYGETGGNSPVQEFTVPYSSSTATISGLSPGVDYTITVYAWGEDSAGYMFMYSPISINYRTC*

AsLOV2-HA4 (C-terminus):
GSSVSSVPTKLEVVAATPTSLLISWDAPMSSSSVYYYRITYGETGGNSPVQEFTVPYSSSTATISGLSPGVDYTITVYAWGEDSAGYMFMYSPISINYRTCGLERIEKNFVITDPRLPDNPIIFASDSFLQLTEYSREEILGRNCRFLQGPETDRATVRKIRDAIDNQTEVTVQLINYTKSGKKFWNLFHLQPMRDQKGDVQYFIGVQLDGTEHVRDAAEREGVMLIKKTAENIDEAAG*

HA4-AsLOV2 (SS59 insertion):
GSSVSSVPTKLEVVAATPTSLLISWDAPMSSSSVYYYRITYGETGGNSPVQEFTVPYSSGLERIEKNFVITDPRLPDNPIIFASDSFLQLTEYSREEILGRNCRFLQGPETDRATVRKIRDAIDNQTEVTVQLINYTKSGKKFWNLFHLQPMRDQKGDVQYFIGVQLDGTEHVRDAAEREGVMLIKKTAENIDEAAGSTATISGLSPGVDYTITVYAWGEDSAGYMFMYSPISINYRTC*

HA4-AsLOV2 (MY90 insertion):
GSSVSSVPTKLEVVAATPTSLLISWDAPMSSSSVYYYRITYGETGGNSPVQEFTVPYSSSTATISGLSPGVDYTITVYAWGEDSAGYMFMGLERIEKNFVITDPRLPDNPIIFASDSFLQLTEYSREEILGRNCRFLQGPETDRATVRKIRDAIDNQTEVTVQLINYTKSGKKFWNLFHLQPMRDQKGDVQYFIGVQLDGTEHVRDAAEREGVMLIKKTAENIDEAAGYSPISINYRTC*

$His_6$-HA4-AsLOV2 (SS58 insertion):
HHHHHHSSGENLYFQGHASGSSVSSVPTKLEVVAATPTSLLISWDAPMSSSSVYYYRITYGETGGNSPVQEFTVPYSGLERIEKNFVITDPRLPDNPIIFASDSFLQLTEYSREEILGRNCRFLQGPETDRATVRKIRDAIDNQTEVTVQLINYTKSGKKFWNLFHLQPMRDQKGDVQYFIGVQLDGTEHVRDAAEREGVMLIKKTAENIDEAAGSSTATISGLSPGVDYTITVYAWGEDSAGYMFMYSPISINYRTC*

HA4-AsLOV2 (YS57 insertion):
GSSVSSVPTKLEVVAATPTSLLISWDAPMSSSSVYYYRITYGETGGNSPVQEFTVPYGLERIEKNFVITDPRLPDNPIIFASDSFLQLTEYSREEILGRNCRFLQGPETDRATVRKIRDAIDNQTEVTVQLINYTKSGKKFWNLFHLQPMRDQKGDVQYFIGVQLDGTEHVRDAAEREGVMLIKKTAENIDEAAGSSSTATISGLSPGVDYTITVYAWGEDSAGYMFMYSPISINYRTC*

His$_6$-HA4-AsLOV2 (SS58 insertion, C450V mutant):
HHHHHHSSGENLYFQGHASGSSVSSVPTKLEVVAATPTSLLISWDAPMSSSSVYYYRITY
GETGGNSPVQEFTVPYSGLERIEKNFVITDPRLPDNPIIFASDSFLQLTEYSREEILGRNVR
FLQGPETDRATVRKIRDAIDNQTEVTVQLINYTKSGKKFWNLFHLQPMRDQKGDVQYFI
GVQLDGTEHVRDAAEREGVMLIKKTAENIDEAAGSSTATISGLSPGVDYTITVYAWGED
SAGYMFMYSPISINYRTC*

SH2-PDC1:
MGSLEKHSWYHGPVSRNAAEYLLSSGINGSFLVRESESSPGQRSISLRYEGRVYHYRINT
ASDGKLYVSSESRFNTLAELVHHHSTVADGLITTLHYPAPKRNKPTVYGVSPNYASSEIT
LGKYLFERLKQVNVNTVFGLPGDFNLSLLDKIYEVEGMRWAGNANELNAAYAADGYA
RIKGMSCIITTFGVGELSALNGIAGSYAEHVGVLHVVGVPSISAQAKQLLLHHTLGNGDF
TVFHRMSANISETTAMITDIATAPAEIDRCIRTTYVTQRPVYLGLPANLVDLNVPAKLLQ
TPIDMSLKPNDAESEKEVIDTILALVKDAKNPVILADACCSRHDVKAETKKLIDLTQFPA
FVTPMGKGSIDEQHPRYGGVYVGTLSKPEVKEAVESADLILSVGALLSDFNTGSFSYSY
KTKNIVEFHSDHMKIRNATFPGVQMKFVLQKLLTTIADAAKGYKPVAVPARTPANAAV
PASTPLKQEWMWNQLGNFLQEGDVVIAETGTSAFGINQTTFPNNTYGISQVLWGSIGFT
TGATLGAAFAAEEIDPKKRVILFIGDGSLQLTVQEISTMIRWGLKPYLFVLNNDGYTIEKL
IHGPKAQYNEIQGWDHLSLLPTFGAKDYETHRVATTGEWDKLTQDKSFNDNSKIRMIEI
MLPVFDAPQNLVEQAKLTAATNAKQ*

His$_6$-SUMO-HA4-AsLOV2 (SS58 insertion, V416I, G528A, N538E mutant):
HHHHHHGSGSGSDQEAKPSTEDLGDKKEGEYIKLKVIGQDSSEIHFKVKMTTHLKKLKE
SYCQRQGVPMNSLRFLFEGQRIADNHTPKELGMEEEDVIEVYQEQTGGHMASKGSSVSS
VPTKLEVVAATPTSLLISWDAPMSSSSVYYYRITYGETGGNSPVQEFTVPYSGLERIEKN
FIITDPRLPDNPIIFASDSFLQLTEYSREEILGRNCRFLQGPETDRATVRKIRDAIDNQTEVT
VQLINYTKSGKKFWNLFHLQPMRDQKGDVQYFIGVQLDGTEHVRDAAEREAVMLIKK
TAEEIDEAAGSSTATISGLSPGVDYTITVYAWGEDSAGYMFMYSPISINYRTC*

His$_6$-HA4-AsLOV2 (SS58 insertion, V416I mutant):
HHHHHHSSGENLYFQGHASGSSVSSVPTKLEVVAATPTSLLISWDAPMSSSSVYYYRITY
GETGGNSPVQEFTVPYSGLERIEKNFIITDPRLPDNPIIFASDSFLQLTEYSREEILGRNCRF
LQGPETDRATVRKIRDAIDNQTEVTVQLINYTKSGKKFWNLFHLQPMRDQKGDVQYFIG
VQLDGTEHVRDAAEREGVMLIKKTAENIDEAAGSSTATISGLSPGVDYTITVYAWGEDS
AGYMFMYSPISINYRTC*

HA4-irFP
MGSSVSSVPTKLEVVAATPTSLLISWDAPMSSSSVYYYRITYGETGGNSPVQEFTVPYSS
STATISGLSPGVDYTITVYAWGEDSAGYMFMYSPISINYRTCGGGAEGSVARQPDLLTC
DDEPIHIPGAIQPHGLLLALAADMTIVAGSDNLPELTGLAIGALIGRSAADVFDSETHNRL
TIALAEPGAAVGAPITVGFTMRKDAGFIGSWHRHDQLIFLELEPPQRDVAEPQAFFRRTN
SAIRRLQAAETLESACAAAAQEVRKITGFDRVMIYRFASDFSGEVIAEDRCAEVESKLGL
HYPASTVPAQARRLYTINPVRIIPDINYRPVPVTPDLNPVTGRPIDLSFAILRSVSPVHLEF
MRNIGMHGTMSISILRGERLWGLIVCHHRTPYYVDLDGRQACELVAQVLAWQIGVMEE
AAATPTCNMRD*

HA4-AsLOV2-irFP (SS58 insertion)
MGSSVSSVPTKLEVVAATPTSLLISWDAPMSSSSVYYYRITYGETGGNSPVQEFTVPYSG
LERIEKNFVITDPRLPDNPIIFASDSFLQLTEYSREEILGRNCRFLQGPETDRATVRKIRDAI
DNQTEVTVQLINYTKSGKKFWNLFHLQPMRDQKGDVQYFIGVQLDGTEHVRDAAERE
GVMLIKKTAENIDEAAGSSTATISGLSPGVDYTITVYAWGEDSAGYMFMYSPISINYRTC
GGGAEGSVARQPDLLTCDDEPIHIPGAIQPHGLLLALAADMTIVAGSDNLPELTGLAIGA
LIGRSAADVFDSETHNRLTIALAEPGAAVGAPITVGFTMRKDAGFIGSWHRHDQLIFLEL
EPPQRDVAEPQAFFRRTNSAIRRLQAAETLESACAAAAQEVRKITGFDRVMIYRFASDFS
GEVIAEDRCAEVESKLGLHYPASTVPAQARRLYTINPVRIIPDINYRPVPVTPDLNPVTGR
PIDLSFAILRSVSPVHLEFMRNIGMHGTMSISILRGERLWGLIVCHHRTPYYVDLDGRQA
CELVAQVLAWQIGVMEEAAATPTCNMRD*

SH2-mCherry-CAAX
MSLEKHSWYHGPVSRNAAEYLLSSGINGSFLVRESESSPGQRSISLRYEGRVYHYRINTA
SDGKLYVSSESRFNTLAELVHHHSTVADGLITTLHYPAPKRNKPTVYGVSPNYVSKGEE
DNMAIIKEFMRFKVHMEGSVNGHEFEIEGEGEGRPYEGTQTAKLKVTKGGPLPFAWDIL
SPQFMYGSKAYVKHPADIPDYLKLSFPEGFKWERVMNFEDGGVVTVTQDSSLQDGEFI
YKVKLRGTNFPSDGPVMQKKTMGWEASSERMYPEDGALKGEIKQRLKLKDGGHYDAE
VKTTYKAKKPVQLPGAYNVNIKLDITSHNEDYTIVEQYERAEGRHSTGGMDELYKGSG
SGSKKKKKKSKTKCVIM

**Supplementary Table 1. Positions within HA4 in which the AsLOV2 domain was inserted.**

| Insertion Name | Insertion Description |
|---|---|
| SS30 | Inserted within S30 and S31 |
| NS47 | Inserted within N47 and S48 |
| SA84 | Inserted within S84 and A85 |
| TG44 | Inserted within T44 and G45 |
| SS58 | Inserted within S58 and S59 |
| SG65 | Inserted within S65 and G66 |
| DA26 | Inserted within D26 and A27 |
| MS29 | Inserted within M29 and S30 |
| MS29-3 | Inserted within M29 and S33 residues 30-32 were removed |
| ED82 | Inserted within E82 and D83 |
| DS83 | Inserted within D83 and S84 |
| GG45 | Inserted within G45 and G46 |
| GN45 | Inserted within G45 and N48 residues 46-47 were removed |
| TN44 | Inserted within T44 and N48 residues 45-47 were removed |
| PT18 | Inserted within P18 and T19 |
| SP68 | Inserted within S68 and P69 |
| SP68-1 | Inserted within S68 and P69 residue 67 was removed |
| N-terminus | Inserted at N-terminus |
| C-terminus | Inserted at C-terminus |
| SS59 | Inserted within S59 and S60 |
| PT18 | Inserted within P18 and T19 residue 17 was removed |
| MY90 | Inserted within M90 and Y91 |
| YS57 | Inserted within Y57 and S58 |

**Supplementary Table 2. Rate and dissociation constants from BLI experiments.**

| Variant | State measured | Illumination | $k_{on}$ ($\mu M^{-1}$ $s^{-1}$) | $k_{off}$ ($s^{-1}$) | $K_d$ ($\mu M$) |
|---|---|---|---|---|---|
| αSH2 Monobody HA4 | - | none | 0.0631 | 0.0145 | 0.23 |
| αSH2-OptoMB$_{C450V}$ | Dark conformation | none | 0.103 | 0.00753 | 0.07 |
| αSH2-OptoMB$_{V416L}$ | Light conformation | 450 nm light | 0.00654 | 0.0733 | 11.2 |
| #αSH2-OptoM$_{V416I_G528A_N538E}$ | Light conformation | not needed* | 0.00225 | 0.0489 | 21.7 |

*Due to the sensitivity of this construct, ambient light in the laboratory as well as internal light from the digital panels of the Octet was sufficient to trigger the lit conformation.

#SUMO tagged

**Supplementary Table 3. Purification parameters of LCAC batch experiments**

| Purified protein | OptoMB version | Resin type | Purity# (%) | Resin Capacity# (mg/mL) | Yield# (% of recovery from the resin) |
|---|---|---|---|---|---|
| YFP-SH2 | Wt AsLOV2 | Talon* | 98.25 (± 2.26) | 5.85 (± 0.67) | 17.8 (± 2.18) |
| YFP-SH2 | #OptoM$_{V416I_G528A_N538E}$ | Talon* | 95.32 (± 0.46) | 4.5 (± 0.33) | 29.79 (± 3.28) |
| SH2-PDC1 | #OptoM$_{V416I_G528A_N538E}$ | Talon* | 95.53 (± 1.05) | 5.24 (± 0.52) | 39.01 (± 2.61) |
| YFP-SH2 | #OptoM$_{V416I_G528A_N538E}$ | CNBr** | 99.78 (± 0.22) | 2.55 (± 0.22) | 40.29 (± 3.02) |
| SH2-PDC1 | #OptoM$_{V416I_G528A_N538E}$ | CNBr** | 96.69 (±1.24) | 1.55 (± 0.18) | 42.28 (± 4.34) |

* OptoMB was His-tag bound

** OptoMB was covalently conjugated

#Calculations are explained in Methods

#SUMO tagged

**Supplementary Table 4. Constructs used in this study.**

| Plasmid | Tag | Protein | Marker | Vector type |
|---|---|---|---|---|
| EZ-L663 | None | HA4 | Kanamycin | pCri-7b |
| EZ-L664 | His$_6$ | YFP-SH2 | Kanamycin | pCri-8b |
| EZ-L703 | None | YFP-SH2 | Kanamycin | pCri-7b |
| EZ-L704 | His$_6$ | HA4 | Kanamycin | pCri-8b |
| EZ-L706 | None | HA4-AsLOV2 (SS58 insertion) | Kanamycin | pCri-7b |
| EZ-L736 | None | HA4-AsLOV2 (MS29 insertion, residues 30-32 were removed) | Kanamycin | pCri-7b |
| EZ-L747 | None | HA4-AsLOV2 (SS59 insertion) | Kanamycin | pCri-7b |
| EZ-L765 | His$_6$ | HA4-AsLOV2 (SS58 insertion) | Kanamycin | pCri-8b |
| EZ-L830 | His$_6$-SUMO | HA4-AsLOV2 (SS58 insertion, V416L mutant) | Kanamycin | pCri-11b |
| EZ-L884 | His$_6$ | HA4-AsLOV2 (SS58 insertion, C450V mutant) | Kanamycin | pCri-8b |
| EZ-L886 | None | SH2-PDC1 | Kanamycin | pCri-7b |
| EZ-L889 | His$_6$-SUMO | HA4-AsLOV2 (SS58 insertion, V416I, G528A, N538E mutant) | Kanamycin | pCri-11b |
| EZ-L892 | His$_6$ | HA4-AsLOV2 (SS58 insertion, V416I mutant) | Kanamycin | pCri-8b |
| AG-pHR1 | None | HA4-iRFP | Ampicillin | pHR |
| AG-pHR2 | None | HA4-AsLOV2-iRFP (SS58 insertion) | Ampicillin | pHR |
| AG-pHR3 | CAAX | SH2-mCherry-CAAX | Ampicillin | pHR |

### Supplementary Figures

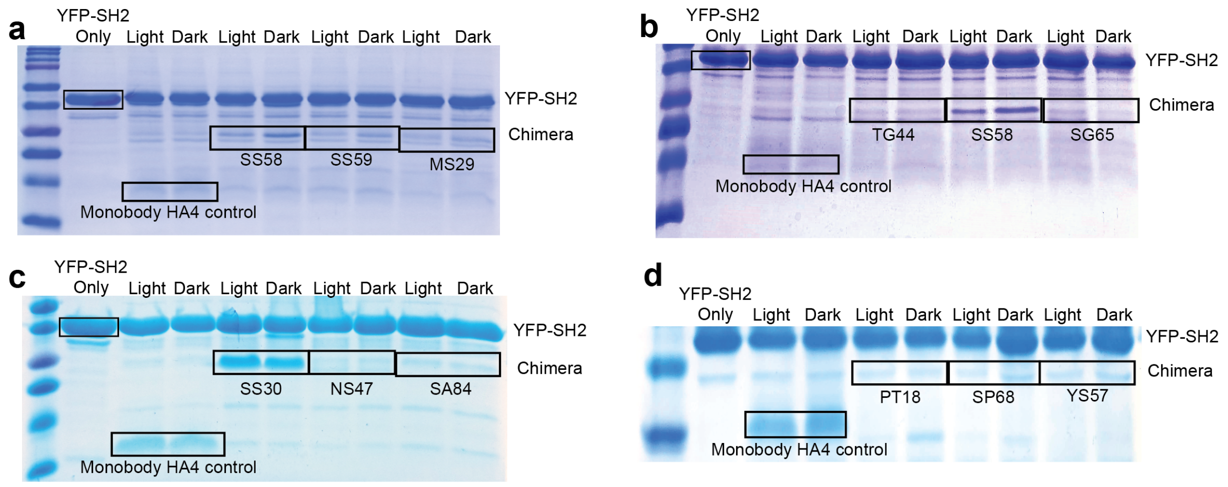


**Supplementary Fig. 1: Representative SDS-PAGE gels of pull-down screens of HA4-AsLOV2 chimeras. a**, Complete SDS-PAGE gel of the results shown in Fig. 1d, including chimeras with the AsLOV2 domain inserted in positions SS58, SS59 and MS29 (with residues S30 to S32 of loop L2 deleted) of HA4. **b-d**, A representative sample of SDS-PAGE gels of other chimeras with the AsLOV2 domain inserted in different positions that either do not bind or bind poorly to SH2 (TG44, NS47, SA84, SG65, SP68, YS57 and PT18); or bind well to SH2, but show no difference in binding between light conditions (SS30).

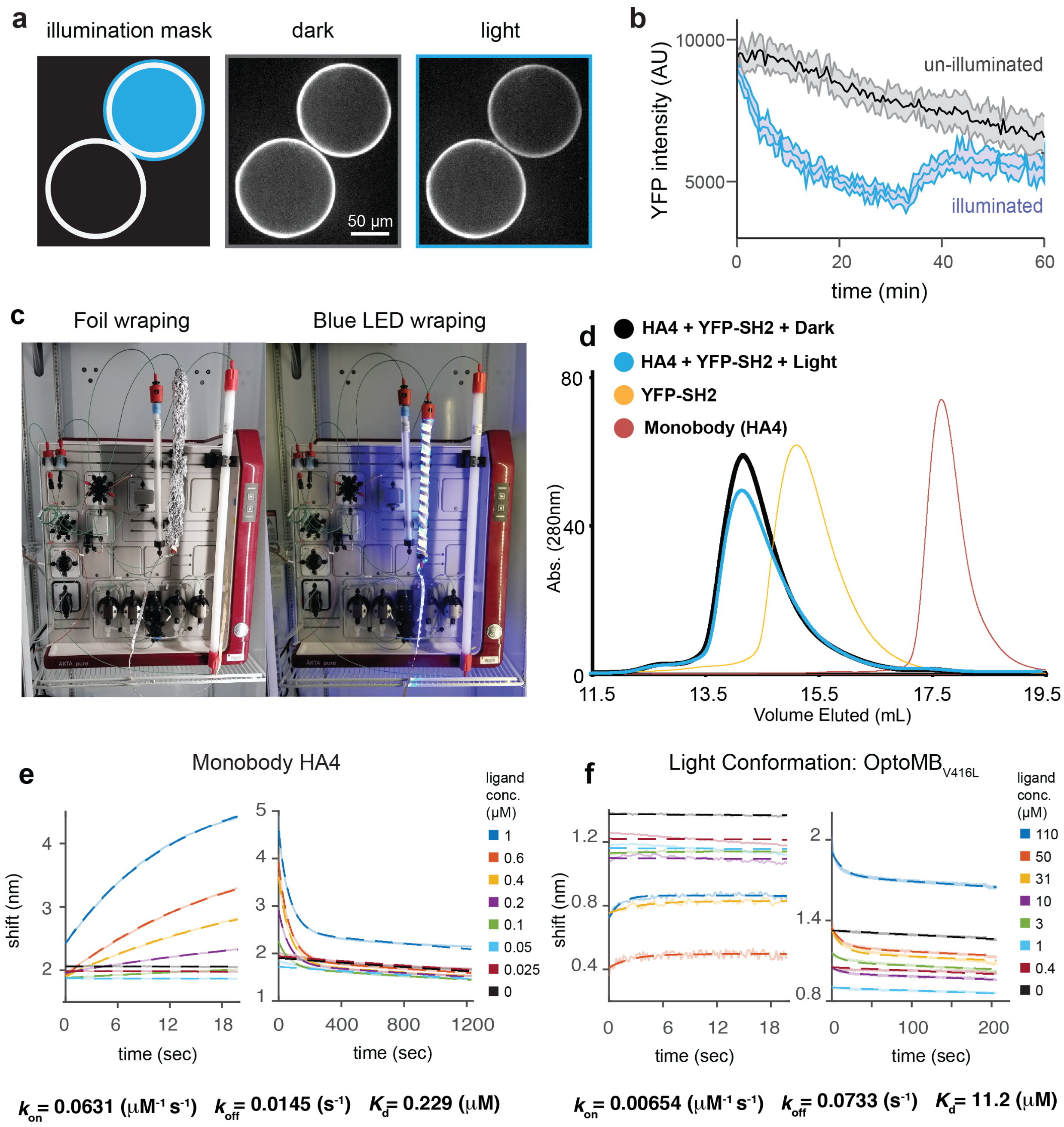


**Supplementary Fig. 2: *In vitro* characterization of OptoMB**. **a**, Light-enabled spatial control of OptoMB binding interactions shown with two OptoMB-coated agarose beads incubated in a circulating solution of YFP-SH2. The blue circle in the left panel indicates the light masked area which was used to illuminated only one bead. The central and right panels show the YFP-SH2 fluorescence on the surface of the beads in the dark and light, respectively, displaying a visible reduction in fluorescence of only the illuminated bead. **b**, Quantification of spatially controlled binding experiments of YFP-SH2 to two OptoMB-coated agarose beads (experiments shown in

**a**), showing YFP fluorescence intensity of both illumined (blue) and unilluminated (gray) beads over time. **c**, Experimental setup of size exclusion chromatography runs using a Superdex 200 16/300 column (GE®). For experiments in the dark (left), the whole column was covered with thick aluminum foil, and the chromatography refrigerator covered with a black blanket to avoid light contamination (not shown). For experiments in the light (right), the column was wrapped with blue LEDs. **d**, Size exclusion chromatography experiments show identical elution profiles of the HA4-YFP-SH2 complex in dark (black) or light (blue) conditions. **e**, BLI measurements of binding (left) and unbinding (right) activities of YFP-SH2 to immobilized HA4 in the dark. **f**, BLI measurements of binding (left) and unbinding (right) activities of YFP-SH2 to immobilized OptoMB exposed to light (using AsLOV2 domain mutant V416L equivalent, which stabilizes the lit-state of AsLOV2). The calculated rate and dissociation constants are shown below the BLI data.

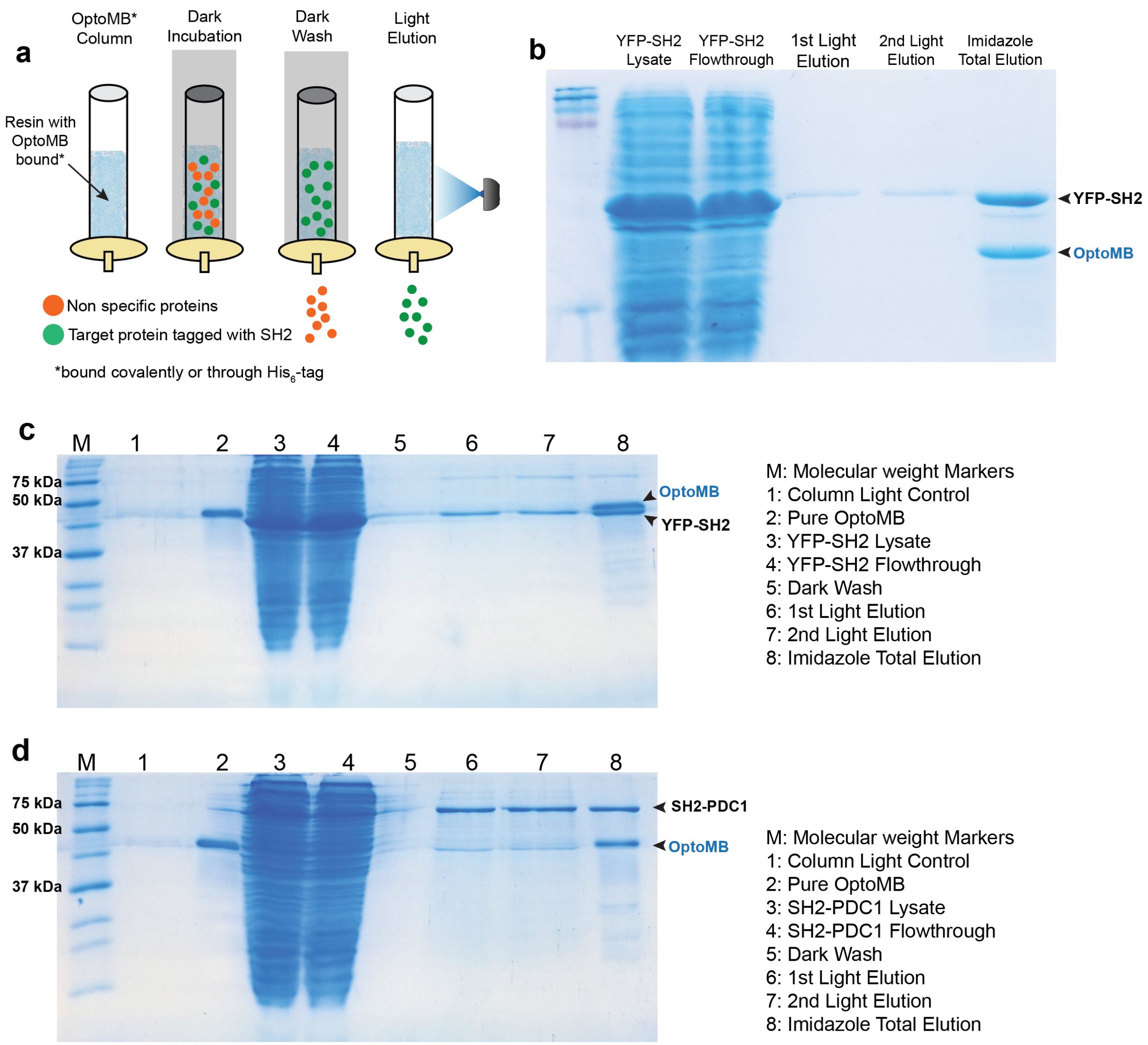


**Supplementary Fig. 3: Light-Controlled Affinity Chromatography (LCAC) to purify SH2-tagged proteins using OptoMB immobilized on $Co^{2+}$ agarose beads. a**, Schematic diagram of LCAC procedure using a column packed with OptoMB-coated agarose beads. After flowing through crude extract and washing in the dark, elution is carried out by applying blue light to the surface of the column. **b**, SDS-PAGE gel of YFP-SH2 purified with OptoMB immobilized column. **c**, SDS-PAGE gel of YFP-SH2 purified using a column packed with agarose beads coated with $His_6$-$OptoMB_{V416I\_G528A\_N538E}$ (with a SUMO tag). **d**, SDS-PAGE gel of SH2-PDC1 purified using a column packed with agarose beads coated with $OptoMB_{V416I\_G528A\_N538E}$ (with a SUMO tag).

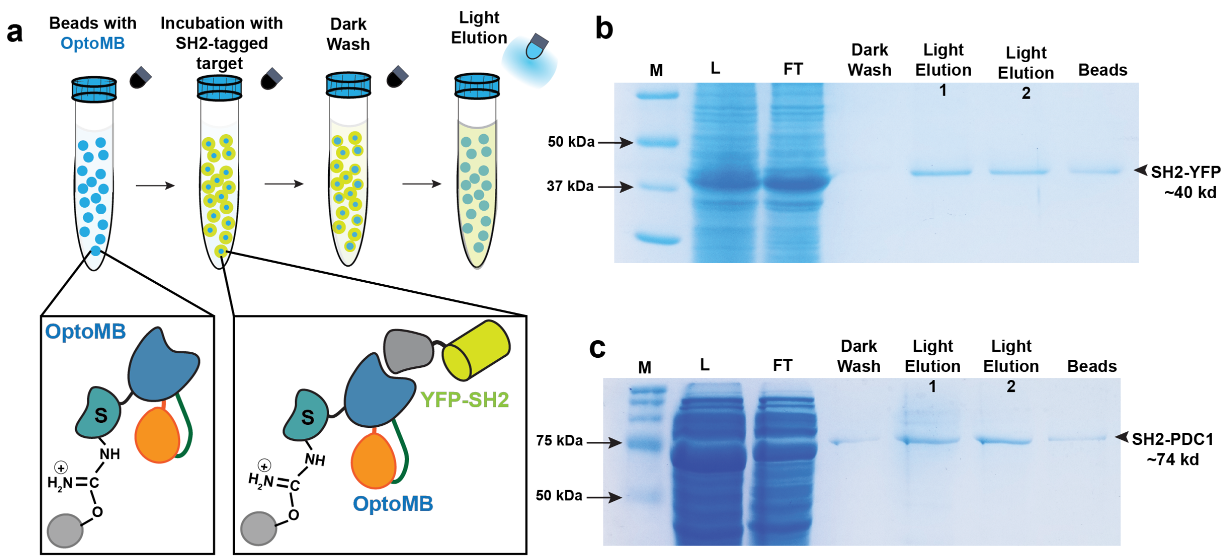


**Supplementary Fig. 4: Light-controlled affinity chromatography to purify SH2-tagged proteins using cyanogen bromide (CNBr)-conjugated OptoMB. a**, Schematic diagram of OptoMB$_{V416I_G528A_N538E}$ with SUMO tag (S) covalently conjugated to CNBr beads through surface-exposed primary amines, and their use in LCAC as described before. **b**, **c**, SDS-PAGE gel of YFP-SH2 (b) or SH2-PDC1 (c) purified using the SUMO-tagged OptoMB$_{V416I_G528A_N538E}$ conjugated to CNBr beads, in batch. Molecular weight markers (M), lysate (L), Unbound flow through (FT), washing step in the dark (Dark Wash), two consecutives light elution aliquots (Light Elution 1 and 2) and heat-treated beads resolved in SDS-PAGE gel (12% polyacrylamide).

**Supplementary Movie legends:**

**Supplementary Movie 1: Modeled Structure of the dark conformation of OptoMB.** Rotation of the energy-minimized structural model of OptoMB shown in Figure 1e. The light-responsive chimera of AsLOV2 (orange), containing the Jα helix that undergoes structural rearrangement upon light stimulation (green), is inserted at position SS58 (in loop L4) of HA4 monobody (blue), bound to SH2 domain (gray). Loop L4, in the monobody, connects βSh2 (blue) with βSh1 (black) forming the core of the monobody fold. Molecules are rotated 720 degrees in both y and x axis.

**Supplementary Movie 2: Time-lapse imaging of Ni-NTA agarose beads coated with His-tagged OptoMB light sensitive variant (V416L) or the control Monobody HA4**. OptoMB coated bead (left) or control Monobody HA4 coated bead (right) were incubated with 2 μM of purified YFP-SH2 in solution (see Methods). Beads were imaged every 2 min using a 20X air objective and a blue LED light (450 nm) was turn on (indicated with the blue box) after beads were equilibrated in the dark. Time is indicated in mm:ss and the scale bar represents 50 μm.

**Supplementary Movie 3: Time-lapse imaging of two Ni-NTA agarose beads coated with His-tagged OptoMB (variant V416L) with spatial illumination of one bead.** OptoMB coated beads were incubated with 2 μM YFP-SH2 in solution (see Methods). Imaging was done every 2 min using a 20X air objective. A masked circular region of blue light (with radius of 70 μm) was set to illuminate only the top right bead and the blue LED light (450 nm) was toggled on and off (indicated with the blue circle in the top right corner). Time is indicated in mm:ss and the scale bar represents 50 μm.

**Supplementary Movie 4: Time-lapse imaging of HEK293T cells expressing OptoMB-irFP (shown) and membrane-localized SH2-mCherry-CAAX (not shown)**. Cells were imaged every 30 sec for 60 minutes using a 60X oil objective. Blue LED illumination (450 nm) was toggled on and off (as indicated by the blue box in the lower left corner). Time is indicated in mm:ss and the scale bar represents 10 μm.
